# A synthetic peptide CTL vaccine targeting nucleocapsid confers protection from SARS-CoV-2 challenge in rhesus macaques

**DOI:** 10.1101/2021.04.24.441228

**Authors:** Paul E. Harris, Trevor Brasel, Christopher Massey, C. V. Herst, Scott Burkholz, Peter Lloyd, Tikoes Blankenberg, Thomas M. Bey, Richard Carback, Thomas Hodge, Serban Ciotlos, Lu Wang, Jason E. Comer, Reid Rubsamen

**Affiliations:** Department of Medicine, Columbia University, P&S 10-502, 650 West 168th Street, New York, NY, USA 10032;.; Department of Microbiology & Immunology, University of Texas Medical Branch, 301 University Blvd., Galveston, TX, 77555, USA; (T.Br.); (J.C.); (C.M.); Flow Pharma Inc., 4829 Galaxy Parkway, Suite K, Warrensville Heights, OH 44128, 94523, USA; (C.H.); (S.B.); (R.C.); (T.H.); (S.C.); (L.W.); (P.L.), (R.R.); Dignity Health Mercy Medical Center, Redding, CA, USA, 96001; (T.Be.); (T.Bl.); The Department of Anesthesiology and Perioperative Medicine, University Hospitals, Cleveland Medical Center, Case Western Reserve School of Medicine, Cleveland, OH, USA; Department of Anesthesia, Critical Care and Pain Medicine, Massachusetts General Hospital, Boston, MA, USA 02114.

**Keywords:** SARS-CoV-2, animal model, macaque, vaccine, MHC Class I peptide, T-cell

## Abstract

**Background:** Persistent transmission of severe acute respiratory syndrome coronavirus 2 (SARS-CoV-2) has given rise to a COVID-19 pandemic. Several vaccines, evoking protective spike antibody responses, conceived in 2020, are being deployed in mass public health vaccination programs. Recent data suggests, however, that as sequence variation in the spike genome accumulates, some vaccines may lose efficacy.

**Methods:** Using a macaque model of SARS-CoV-2 infection, we tested the efficacy of a peptide-based vaccine targeting MHC Class I epitopes on the SARS-CoV-2 nucleocapsid protein. We administered biodegradable microspheres with synthetic peptides and adjuvants to rhesus macaques. Unvaccinated control and vaccinated macaques were challenged with 1 x 10^8^ TCID_50_ units of SARS-CoV-2, followed by assessment of clinical symptoms, viral load, chest radiographs, sampling of peripheral blood and bronchoalveolar lavage (BAL) fluid for downstream analysis.

**Results:** Vaccinated animals were free of pneumonia-like infiltrates characteristic of SARS-CoV-2 infection and presented with lower viral loads relative to controls. Gene expression in cells collected from BAL samples of vaccinated macaques revealed a unique signature associated with enhanced development of adaptive immune responses relative to control macaques.

**Conclusions:** We demonstrate that a room temperature stable peptide vaccine based on known immunogenic HLA Class I bound CTL epitopes from the nucleocapsid protein can provide protection against SARS-CoV-2 infection in non-human primates.

**Graphical Abstract:** 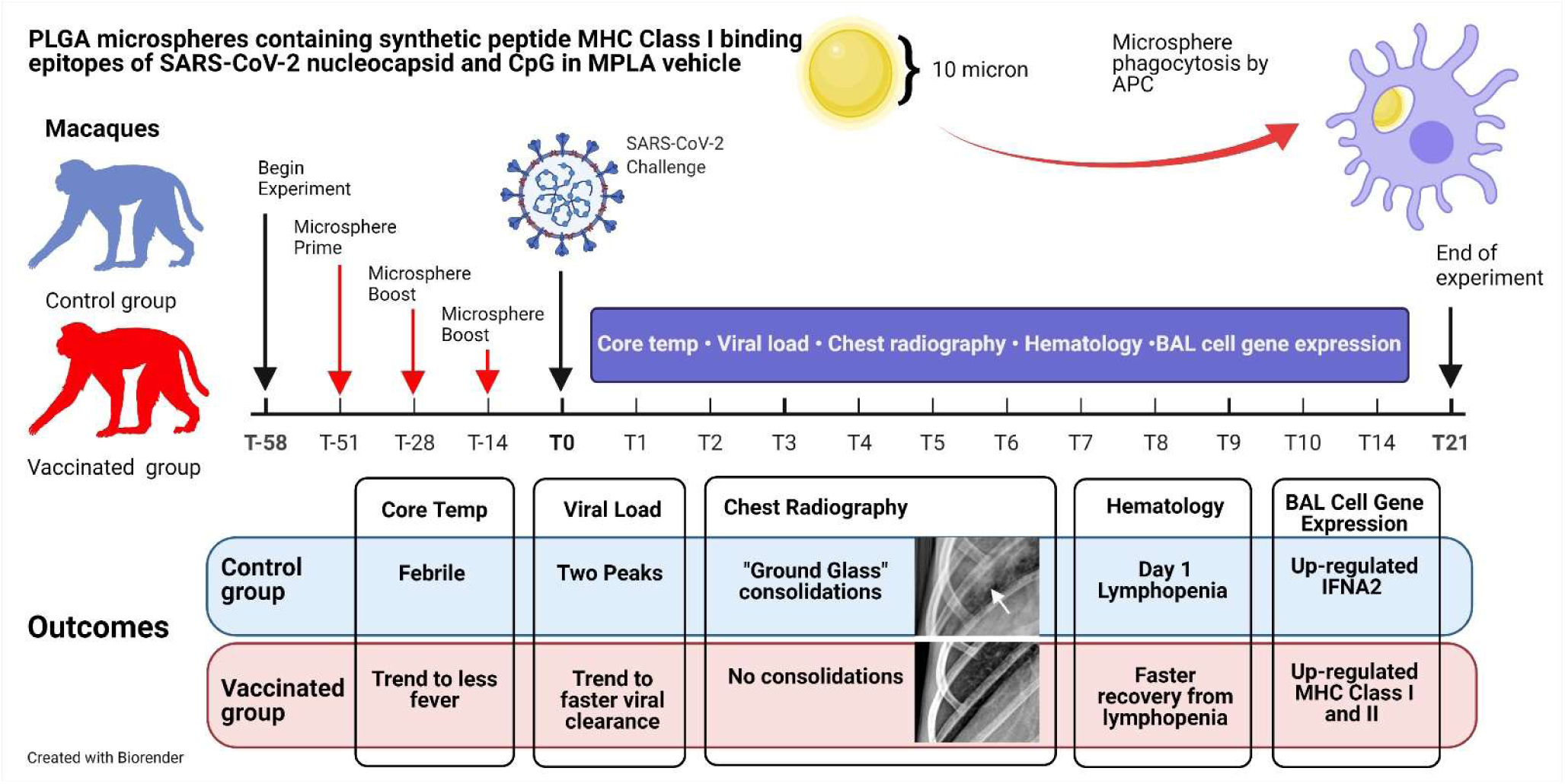

## 1. Introduction

As of April 2021, over 1.6% of the world population have had confirmed COVID-19 disease and the new case rate is about one-half million per day. Less than 2 % of the world’s population are vaccinated against SARS-CoV-2[1]. Confounding efforts to reach herd immunity to COVID-19 disease include, but are not limited to the following: 1) the spread of SARS-CoV-2 mutations affecting the efficacy of current iterations of vaccines and therapeutic biologics[2], 2) the speed of SARS-CoV-2 vaccine deployment, development, and manufacture, and 3) in the context of global public health, issues related to vaccine hesitancy, cold supply chain requirements, the total manufacturing cost per dose, and ease of administration.

We have previously described a novel vaccine platform [3, 4], which may address many of the above concerns and now report on its efficacy in a rhesus macaque model of SARS-CoV-2 infection. Macaque models of COVID-19 disease have been previously reported and have served as critical tools for understanding disease pathology and for the development and testing of vaccines and therapeutics [5–21]. While the clinical course of SARS-CoV-2 infection in macaques is milder relative to that observed in humans [6,8,20,21], the macaque model remains the gold standard for preclinical evaluation of COVID-19 vaccines [10,19,22–27]. Our overall approach focuses on promoting protective T-cell immunity using synthetic peptides delivered in biodegradable microspheres together with Toll-like receptor (TLR) 4 and 9 adjuvants and differs from current COVID-19 vaccines against spike proteins. The synthetic peptide sequences applied are based on known immunogenic HLA Class I bound epitopes that have previously been characterized in either SARS-CoV-1 [28] or SARS-CoV-2 [29–31] infections. We provide evidence that application of this vaccine platform in SARS-CoV-2 challenged macaques provides protection from pneumonia-like pathology observed in virally challenged but unvaccinated control non-human primate (NHP) subjects, reduces viral loads as compared to unvaccinated controls, and induces changes in the gene expression patterns in recovered BAL cells consistent with enhanced antigen presentation capacity and markers of T-cells.

## 2. Materials and Methods

### A. Macaque MHC Class I Typing

The MHC Class I genes of a cohort of 15 rhesus macaques (Envigo, Alice, TX, USA) were molecularly-typed by the University of Wisconsin-Madison National Primate Research Center. Mamu (*Macaca mulatta*) MHC class I alleles were typed amplifying the genomic DNA of each subject using a panel of specific primers for exon 2 of all known MHC-A and MHC-B alleles encoded by each subject’s target DNA. Resulting amplicons were sequenced by the Illumina MiSeq method [32]. The primer panel contained specific primer pairs able to amplify all possible MHC-A and MHC-B alleles encoded by each macaque. Sequencing data analysis provided a high-resolution haplotype for the MHC-A and MHC-B alleles carried by each subject. Following analysis, we selected four of the 15 macaques for vaccination based on peptide MHC binding predictions as described below.

### B. Vaccine Design/Peptide Selection and Manufacture

#### In silico design and selection of SARS-CoV-2 CTL epitopes

The overall strategy and rationale used in the selection of synthetic peptides used to stimulate potential CTL immune responses in SARS-CoV-2 infected humans has been previously described [3, 4]. We selected SARS-CoV-2 nucleoprotein as the target of CTL attack based on the following rationale: 1) survivors of SARS-CoV-1 have shown a memory T-cell response to nucleoprotein at least 2 years after infection [28], 2) there is >90% amino acid sequence homology between SARS-CoV-1 and SARS-CoV-2 nucleoprotein (the homology for the selected CTL epitopes used in this report is 100%) [33], and 3) in general, there is a lower frequency of mutations resulting in amino acid substitutions (relative to Spike protein) that might affect the immunogenicity of the selected CTL epitopes represented by synthetic peptides within the vaccine formulation [34]. The lower mutation frequency may reflect the hypothesis that amino acid substitutions in nucleoprotein may impact viral fitness [35]. We reviewed previous literature and MHC peptide-binding databases [36] and selected five amino sequences representing SARS-CoV-2 nucleoprotein with predicted strong in vitro affinity for HLA Class I molecules[37], and/or documented or predicted immunogenic potential[4,28–31,33,38,39]. Together this set of peptides yielded potential broad coverage of HLA haplotypes (>90% worldwide) (Supplemental data, Table S1A). The predicted binding of this set of peptides was examined within the MHC genotypes of the cohort of 15 rhesus macaques [40] available to us and the best correspondence between selected peptides and rhesus MHC Class I genotype was selected (Supplemental data, Table S1B). Because the predicted peptide macaque MHC binding coverage for the peptide LLLDRLNQL was incomplete in the available genotypes, we added an additional peptide (ASAFFGMSR) with predicted strong Mamu MHC Class I binding to the formulation for a total of six peptides.

#### Microsphere preparation and adjuvant formulation

The peptide epitopes used in this study were delivered in vivo by intratracheal instillation of a formulation of Poly-L-lactide-co-glycolide (PLGA) microspheres containing the corresponding synthetic nine-mer peptides and TLR-9 agonist CpG oligonucleotide adjuvant in a vehicle containing TLR-4 agonist monophosphoryl lipid A (MPLA). The rationale for the choice of the delivery platform and the basic manufacturing scheme used in production has been previously reported [3,4,41]. Briefly, room temperature solutions of a synthetic peptide, CpG oligonucleotide, and mannose were mixed with a solution of PLGA in acetone/water followed by sonication. The formulation was then processed through a precision spray-drying device (Buchi Corporation, New Castle, DE, USA) and passed through a drying chamber (air at room temperature) to allow evaporation of the acetone. The dry microsphere stream was analyzed in real-time through a laser particle size analyzer (SprayTech, Malvern Instruments, Malvern, PA) before collection (Buchi cyclone drier) as a dry powder for reconstitution at the time of delivery using a 2% DMSO aqueous solution containing MPLA (20 μg/ml). Each microsphere contained peptide loaded at approximately 0.1% by weight and CpG 0.01% by weight. Monitoring of the microsphere diameters allowed the production of microspheres with a mean diameter of 10 ± 2 microns. This diameter was selected for formulation to ensure delivery via phagocytosis of no more than 1-4 microspheres per antigen-presenting cell (APC) which have an average diameter of 13 microns [3]. cGMP manufacturing protocols were employed using GMP grade synthetic peptides (Peptides International, Louisville, KY, USA), CpG oligonucleotides (Trilink Biosciences, San Diego, CA, USA), and MPLA (Avanti Polar Lipids, Alabaster, AL, USA). The CpG oligonucleotide and MPLA used in this study were manufactured using the same chemical compositions as equivalent materials used in FDA-approved vaccines. Assessment of thermal stability of the synthetic peptides within the microspheres has been previously reported [3]. Peptide content and structure in microspheres were determined by HPLC after two months of room temperature storage. We found that over 99% of the peptide was maintained structurally intact (data not shown).

### C. Animal Studies

#### Ethics statement

The animal research protocols used in this study were performed in strict accordance with the recommendations in the Guide for Care and Use of Laboratory Animals, Eighth Edition (National Academy Press, Washington, D.C., 2011). The University of Texas Medical Branch (UTMB) facility where these studies were conducted is accredited by the Association for Assessment and Accreditation of Laboratory Animal Care. The protocols were approved by the UTMB Institutional Animal Care and Use Committee (Protocol Numbers 2004051 [natural history/control study] and 2003033 [vaccination study]) and complied with the Animal Welfare Act, the U.S. Public Health Service Policy, and other Federal statutes and regulations related to animals and experiments involving animals. All hands-on manipulations, including immunizations and biosampling, were performed while animals were sedated via ketamine (5 mg/kg)/dexmedetomidine (0.025 mg/kg) intramuscular injection. All efforts were made to minimize suffering.

#### Macaques

Adult Indian origin rhesus macaques (*Macaca mulatta*, n = 7 [5 male, 2 female], 46-48 months old) or Vietnamese origin cynomolgus macaques (*Macaca fascicularis*, n = 1 female, 84 months old), individually identified via unique tattoo, were obtained from Envigo/Covance (Alice, Texas, USA). All animals were considered healthy by a veterinarian before being placed on study. Macaques were individually housed in stainless steel nonhuman primate caging equipped with squeeze backs for the duration of the studies. For continuous core body temperature measurements, a DST micro-T implantable temperature logger (Star–Oddi, Gardabaer, Iceland) was surgically implanted into the peritoneal cavity of each animal prior to study initiation; data recording was set to 10- or 15-min intervals for control and vaccinated macaques, respectively. Certified primate Diet 5048 was provided to the macaques daily. Drinking water (RO) was provided ad libitum through an automatic watering system. To promote and enhance the psychological well-being of the animals, food enrichment consisting of fresh fruits and vegetables was provided daily. Environmental enrichment including various manipulatives (Kong toys, mirrors, and puzzles) was also provided.

### D. Immunization, Virus Challenge, Post-Challenge Monitoring and Biosampling

#### Immunization and ELISPOT analysis

Immunizations were performed on the selected MHC-typed rhesus macaques (n = 4) via ultrasound-guided inguinal lymph node injection (LN) and/or intratracheal instillation (IT). Twenty mg of vaccine microsphere preparation in 1ml was used for each LN injection (two injections / dose / animal) and 100mg of vaccine microspheres in a 5 ml volume was used for each IT administration. Specifically, on Day -51 (51 days prior to virus challenge), two of the macaques (Figure 1) were administered 2 mL of vaccine via LN injection (1 mL per node). Subsequent administration of the vaccine occurred via the IT route (5 mL) only as described previously[42]. Remaining vaccine doses, administered on Days -28 and -14, were delivered via IT only (5 mL per dose) to the rhesus vaccination group. On Days -44, -21, and -7 (7 days post-vaccination), femoral vein peripheral blood (8 mL) was collected from each animal into a BD Vacutainer® CPT^TM^ Cell Preparation Tube with Sodium Heparin (Becton, Dickinson and Company, Franklin Lakes, NJ, USA) and processed to peripheral blood mononuclear cells (PBMCs) per manufacturer instructions. Collected PBMCs were assessed for immunoreactivity via ELISPOT. In brief, ELISPOT assay plates (MabTech Inc., Cincinnati, OH, USA) specific for the detection of primate IFNγ were used according to manufacturer instructions. BAL cell concentrations were adjusted to 1 x 10^5^ cells per mL in a complete growth medium. Diluted BAL cells were dispensed (100µL/well) into a 96-well plate after which 100µL of complete growth medium (CGM, negative control), Concanavalin A in CGM at 10 µg per well (positive control), and various concentrations of specific (i.e., immunizing) and non-specific peptides (Supplemental data, Table S1B) were added. Peptides used for immunization were added to wells at a concentration of 50 µM. All samples were assayed in duplicate. Plates were incubated at 37°C/5% CO_2_ for 20-22 hours after which plates were thoroughly washed. Conjugated detection antibody was then added and incubated followed by additional washing. Wells were developed using TMB as a substrate. Counts were performed at Cellular Technology Corporation (Shaker Heights, OH, USA) using an Immunospot Analyzer and all well images were quality-controlled on site. All spot-forming cell counts reported are the result of averaging counts from the duplicate 50µM immunization-specific peptide wells.

**Figure 1.**
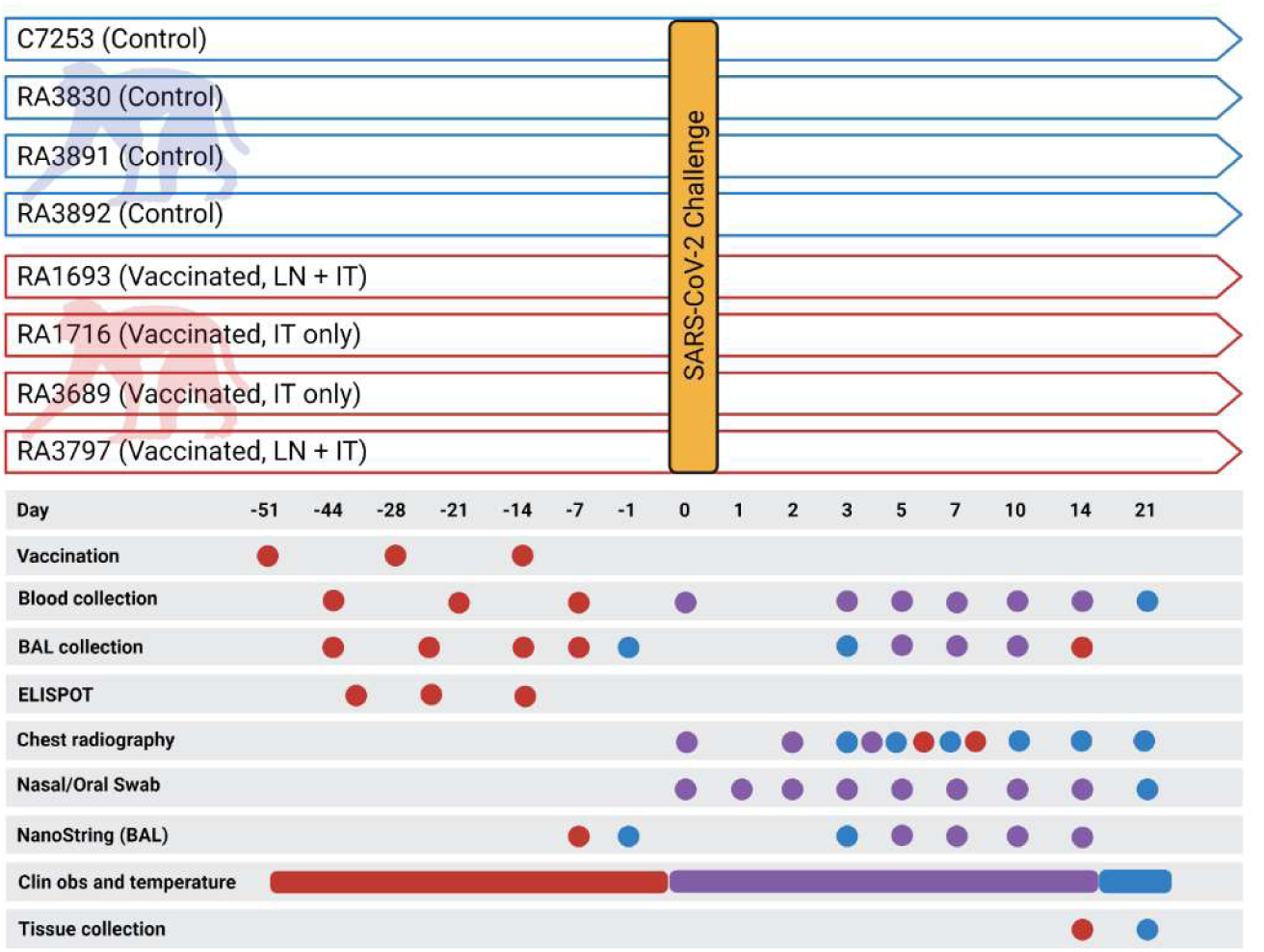
Schematic of the experimental protocol. Unvaccinated (control) macaques are represented by blue coloring. Vaccinated macaques are represented by red coloring. Overlapping tasks are represented by purple coloring. Graphic created with BioRender.com.

#### Virus challenge

On Day 0, macaques were administered 1-5 ×10^8^ TCID_50_ SARS-CoV-2 (USA_WA1/2020) via combined mucosal atomization (1 mL as delivered using a MAD Nasal^TM^ Intranasal Mucosal Atomization Device per manufacturer instructions) and intratracheal instillation (4 mL). Intratracheal instillations were performed as described above for delivery of the vaccine. The virus suspension was prepared on the day of challenge from frozen seed stock (kindly provided by Dr. Chien-Te [Kent] Tseng at UTMB) initially generated (one passage) in Vero C1008 (E6) cells (BEI Resources, NR-596, Lot 3956593) from original material provided by the Centers for Disease Control and Prevention in January 2020. Next-generation sequencing confirmed a 100% consensus sequence-level match to the original patient specimen (GenBank accession MN985325.1).

#### Post-challenge monitoring and chest radiography

Animals were monitored and scored twice daily for clinical signs of disease including alterations in activity/appearance (i.e., hunched posture), food consumption/waste output, and were scored based on general appearance, activity, food consumption, and outward changes in breathing patterns. Prospectively defined criteria that required immediate euthanasia included severe dyspnea and/or agonal breathing and prostate posture/reluctance to move when stimulated. No animals met endpoint criteria during the study. Ventrolateral chest radiography was performed on the days indicated (Figure 1) using a portable GE AMX-4+ computed radiography system per manufacturer instruction. DICOM data files were independently evaluated by two independent investigators blinded to group assignment with large animal imaging experience via a four-pattern approach (analyses of consolidation, interstitial areas, nodules or masses, and atelectasis).

#### Biosampling

Blood, nasal cavity samples, and BAL fluid were collected at the indicated times (Figure 1). Femoral vein peripheral blood was collected via Vacutainer® into standard collection tubes containing ethylenediaminetetraacetic acid (EDTA). Hematology was performed on EDTA blood using the Abaxis VETSCAN® HM5 Hematology Analyzer (Abaxis, Inc., Union City, CA, USA). Nasal cavity samples, collected using sterile cotton-tipped medical swabs, were placed into 0.5 mL sterile phosphate-buffered saline (PBS) for viral load analysis. For BAL fluid collection, animals were sedated as previously described and placed in ventral recumbency. The trachea was visualized and cannulated by an appropriately sized rubber feeding tube. Following the placement of the feeding tube, 20mL of sterile PBS was introduced into the lung and recovered manually through the feeding tube via syringe. This was repeated for a total of 40mL per animal. The total collected volume from each animal (10-30mL) was pooled and centrifuged under ambient conditions (10 min at 500 x g) after which the supernatant was removed. The resulting cell pellet was resuspended in 2 mL of sterile PBS. From this, 1 mL was used for ELISPOT analysis as described for PBMCs. The remaining volume was used for viral load analysis and gene expression profiling.

### E. Viral Load Analysis

#### Infectious viral load (TCID_50_)

Nasal swab and BAL cells suspension samples were serially diluted and incubated with 2×10^4^ Vero C1008 (E6) cells (BEI Resources, NR-596, Lot 3956593) in 100 μl of culture medium (MEM/2% FBS) in 96-well flat-bottom plates (n = 5 replicate wells per dilution). Each plate contained negative and positive control wells inoculated with culture medium and diluted virus stock, respectively. Cultures were incubated at 37°C/5% CO2 for 96h after which cytopathic effect was measured via microscopic observation. The TCID_50_/mL value for each sample was calculated as previously described[43]. Macaque C75243 (cynomolgus) was not included in this study.

#### qRT-PCR

Nasal swab and BAL cell suspension samples (50μL) were added to TRIzol® LS Reagent (250μL) and allowed to incubate under ambient conditions for 10 min. Samples were processed to RNA using Zymo Direct-zol^TM^ RNA Mini Prep kits per manufacturer instructions. RNA samples were analyzed via qRT-PCR targeting the SARS-CoV-2 E gene. Probe (Integrated DNA Technologies, Coralville, IA, USA) was labeled at the 5’-end with fluorophore 9-carboxyfluoroescein (6-FAM) and included an internal quencher (ZEN) and a 3’-end quencher (IowaBlackFQ, IABkFQ). Master Mix was prepared by combining forward primer (250 nM, 5’-ACAGGTACGTTAATAGTTAATAGCGT-3’), reverse primer (250 nM, 5’-ATATTGCAGCAGTACGCACACA-3’), and probe (375 nM, 5’-6FAM-ACACTAGCC/ZEN/ATCCTTACTGCGCTTCG-IABkFQ-3’) with 12.5μL of 2X QuantiFast Probe Mix (QIAGEN), 0.25μL of 2X QuantiFast RT Mix (QIAGEN), and PCR-grade water (fill to 20 μL). To the Master Mix, a test sample (5μL) was added resulting in a final volume of 25μL per reaction. Real-time analysis was performed using the Bio-Rad CFX96^TM^ Real-Time PCR Detection System. Thermocycling conditions were as follows: Step 1, 1 cycle, 50°C for 10 minutes; Step 2, 1 cycle, 95°C for 10 minutes; Steps 3-5, 45 cycles, 95°C for 10 seconds, 60°C for 30 seconds, single read. Negative controls included reaction mixtures without RNA. For quantification purposes, viral RNA extracted from the virus seed stock with a known TCID_50_/mL titer was used. All qRT-PCR results are expressed as TCID_50_/mL equivalents. Macaque C75243 (cynomolgus) was not included in this study.

### F. Gene Expression Profiling

BAL samples were processed to RNA as described above for qRT-PCR analysis. RNA quantity and quality were assessed using a NanoDrop^TM^ Lite Spectrophotometer (ThermoFisher Scientific, Waltham, MA, USA). Samples, normalized to 20ng/μL, were analyzed by NanoString Technologies (Seattle, WA, USA) using the nCounter® SPRINT™ Profiler gene expression profiling using the Non-Human Primate Immunology V2 Panel containing 754 genes that encompass 17 immune-related signaling pathways with isoform coverage for both *Macaca mulatta* and *Macaca fascicularis*. Probe sets that did not cover both *Macaca* species were eliminated resulting in a probe set of 730 genes. Raw gene expression data sets received from NanoString Technologies were processed to remove background signals and normalized using the nSolver^TM^ V.3.0 digital analyzer software. Background signal correction was accomplished by subtracting the NanoString negative control genes. Gene expression normalization was performed using the 16 internal reference genes included in the panel.

### G. Study Termination

At scheduled study termination time points (14 and 21 days post-challenge for vaccinated and control macaques, respectively), animals were humanely euthanized via intravenous administration of a pentobarbital-based euthanasia solution under deep anesthesia followed by bilateral thoracotomy.

### H. Statistical Analysis

Descriptive statistics were performed using Microsoft Excel. Hypothesis testing was performed by considering the null hypothesis of the absence of an association between the compared variables. The statistical strength of associations of continuous data was tested using Students *t*-testing. Qlucore Omics Explorer 3.5 (Qlucore), Metascape (Metascape.org) was used to identify the discriminating variables within the NanoString gene expression data sets from BAL sample analysis that were most significantly different between vaccinated and control subjects. The identification of significantly differential variables between the two groups was performed by fitting a linear model for each variable. The set of genes (87 variable genes out of a total of 730 genes) was identified using a p-value of 0.05, at least a three-fold change, and a q-value cutoff of 0.1. P-values were adjusted for multiple testing using the Benjamini-Hochberg method [44]. Gene expression data were scaled to a mean = 0 and a variance = 1 before clustering. Hierarchical clustering of gene expression in BAL was performed using a supervised weighted average linkage two comparisons approach. The metric used in scaling dendrogram arms was Pearson’s correlation coefficient.

## 3. Results

### A. Primary Clinical Outcomes

#### Clinical signs, body temperature alterations and hematology

Following SARS-CoV-2 challenge, outward clinical signs measured in control macaques included acute mild lethargy and respiratory distress. All vaccinated animals were normal throughout the post-challenge study period. Core body temperatures, as measured via implanted Star-Oddi DST temperature loggers, demonstrated a disruption in the diurnal cycle and mild fever lasting 2-5 days post-challenge in all four-control animals (Figure 2, top panels). Conversely, only two of the four vaccinated macaques (RA1693 and RA3797) presented with similar findings, although diurnal cycle disruption was of shorter duration (1-2 days) and the febrile response was milder (Figure 2, bottom panels). No alterations were measured in the remaining two vaccinated animals. Prior to virus challenge, vaccinated macaques presented with occasional disruptions in the diurnal temperature associated with the vaccination procedure (Supplemental Data, Figure S1).

**Figure 2.**
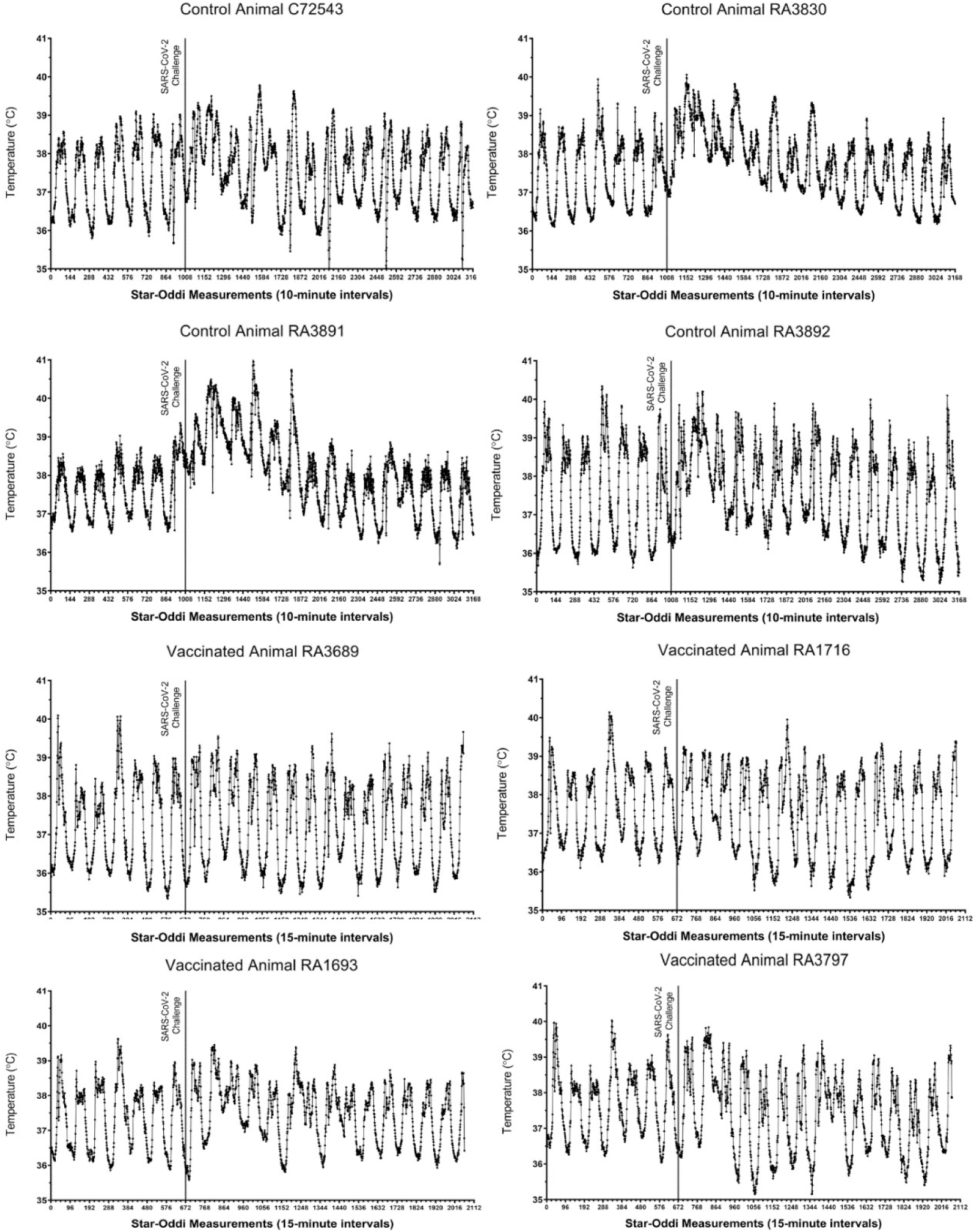
Core body temperature alterations in control and vaccinated macaques following SARS-CoV-2 challenge. For each animal, seven days of pre-challenge baseline temperature measurements are shown. Each tick on the x-axis represents 6 hours or 36 individual logger measurements.

Automated hematology analyses were performed on peripheral blood samples (Supplemental Data, Figure S2). Overall, the number of white blood cells was significantly increased in control subjects on post-challenge days 3 and 5. We observed general lymphopenia in all macaques on day 1 post-challenge. By day 3 post-challenge, lymphocyte counts significantly increased in vaccinated subjects relative to unvaccinated control subjects. Peripheral blood monocyte counts generally peaked on days 1 through 5 in all animals but remained significantly elevated in vaccinated macaques at the end of the study. Neutrophil counts generally rose by day 1 post-challenge in all subjects. There was a transient significant elevation in neutrophils on day 7 post-challenge in vaccinated animals.

#### Viral load

Following SARS-CoV-2 challenge, nasal swab and BAL fluid samples were collected throughout the post-challenge period for analysis of infectious viral load and viral RNA via TCID_50_ and qRT-PCR assays, respectively. Infectious virus was measured from nasal swabs of control and vaccinated macaques beginning one-day post-challenge (Figure 3 [top panel]). By Day 7, 3 of the 4 unvaccinated animals continued to demonstrate infectious viral shedding, albeit at low levels. In contrast, infectious virus could be measured in only one of the vaccinated macaques at the same time point. By 10 days post-challenge, infectious viral loads were undetectable in nasal swab samples from all animals. Viral RNA in nasal swabs generally reflected infectious viral loads. By Day 7, three of the four vaccinated animals demonstrated a 100-fold decrease in nasal swab viral RNA relative to the unvaccinated controls (Figure 3 [bottom panel]). By 14 days post-challenge, SARS-CoV-2 RNA levels were undetectable in nasal swab samples from all subjects. Infectious viral load data were used to calculate an average viral clearance rate post-challenge for each rhesus macaque. The average viral clearance rate from Days 2 through 10 was 4-5 fold higher in two of the four vaccinated macaques (RA1693 and RA3797) relative to the unvaccinated controls (Supplemental data, Figure S3).

**Figure 3.**
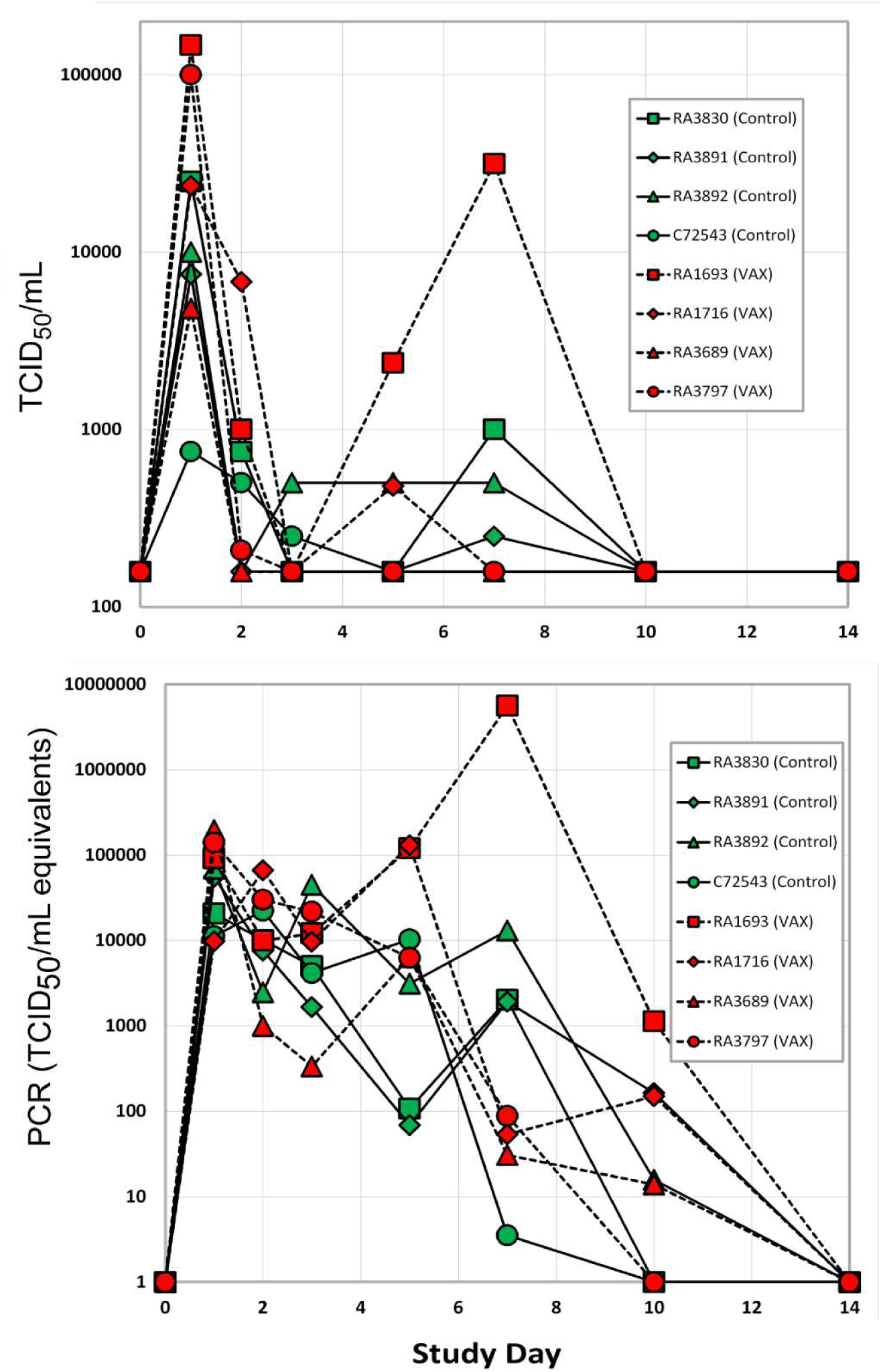
Viral load in nasal swab samples as measured via TCID_50_ assay (top) and qRT-PCR (bottom). The LLOD of the plaque assay was 150 units. Red symbols are vaccinated rhesus macaques, control unvaccinated rhesus subjects are shown in green symbols.

#### Macaque chest radiography

SARS-CoV-2 challenge in unvaccinated controls resulted in mild-to-moderate lung abnormalities, similar to those previously reported for macaques [6,15,16,20,45–48]. These were predominantly limited to the caudal lung relative to baseline images, peaked 3-5 days post-challenge, and were qualitatively characteristic of subclinical or mild-to-moderate human COVID-19 (e.g., ground-glass opacities with or without reticulation, paving, silhouetting, and/or linear opacities). The mild to moderate interstitial pneumonitis seen on the ventrolateral chest radiographs of unvaccinated subjects are consistent with focal infiltrates representing a complex of interstitial macrophages, neutrophils, and plasmacytoid dendritic cells [49]. Abnormalities in control animals resolved by Days 10-21. In contrast, vaccinated macaques lacked the appearance of ground-glass opacities in all regions of the lung throughout the study period (Figure 4 and Supplemental data, Figures S4-S9). We did observe, however, modest bilateral increases in reticulation in vaccinated macaques on Days 3-5, but these abnormalities also resolved by Day 10-21. Bronchoalveolar lavage has been reported to affect computerized tomography X-ray results in healthy rhesus macaques [50]. In our study, however, the pattern of reported changes in vaccinated (i.e., healthy) animals on which BAL was performed was more similar to increases in reticulation versus the patchy consolidations observed in the unvaccinated controls.

**Figure 4.**
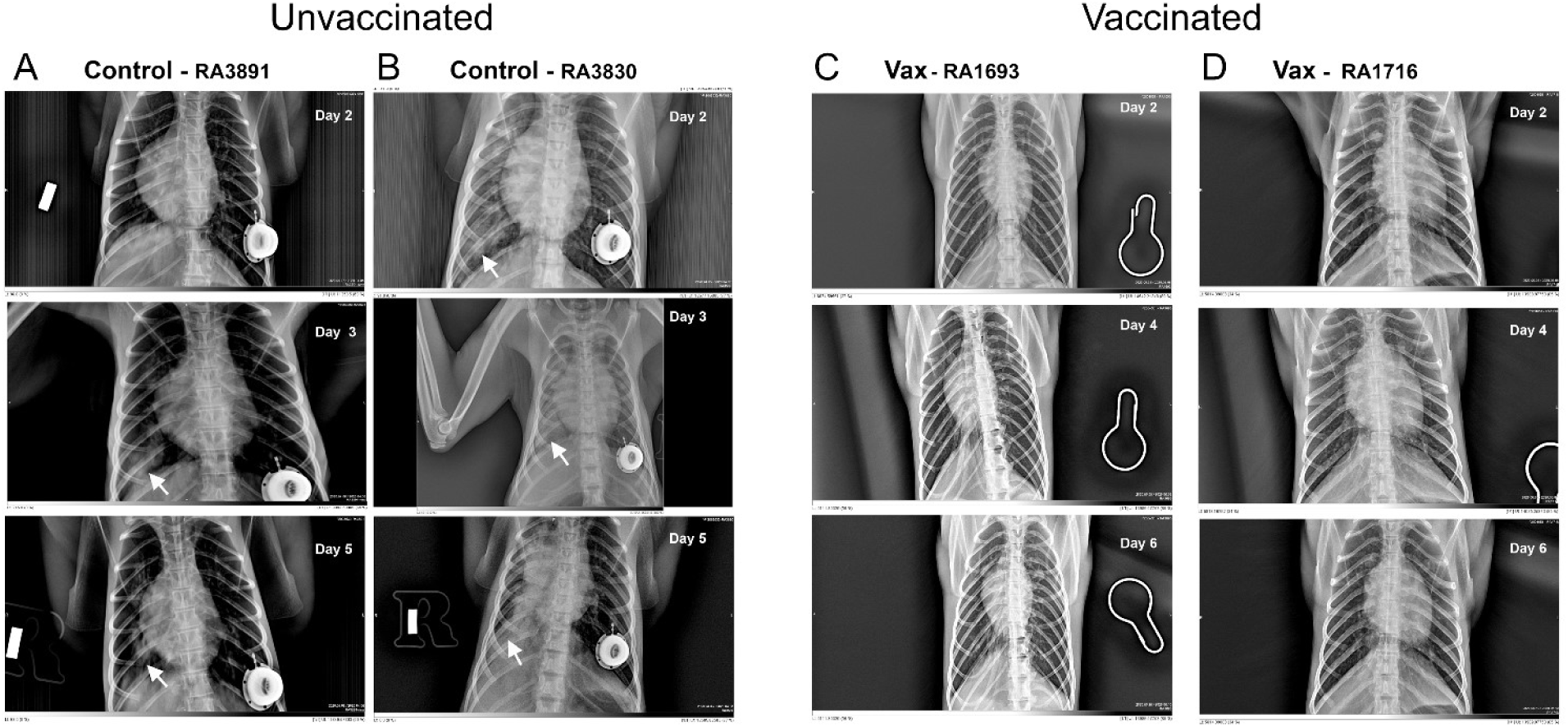
Representative chest radiographs of control and vaccinated macaques following SARS-CoV-2 challenge. As shown, control macaques (left columns A and B) demonstrated a progression of pulmonary infiltrates during the acute period (Days 2-5) of disease post-challenge. In contrast, vaccinated macaques (right columns C and D) lacked similar abnormalities. White arrows indicate areas of mild to moderate pulmonary infiltrates seen as ground glass consolidations.

### B. Secondary Outcomes

#### Analysis of gene expression patterns in BAL cells

We identified a set of 87 genes in BAL samples collected 5-7 days post-challenge from control animals with statistically significant differential expression (as measured from changes in accumulation of their specific transcripts) versus BAL samples collected from vaccinated animals during the same time points (Figures 5 and 6). We selected to focus on the Day 5 and Day 7 samples to capture a possible peak of adaptive immune responses to SARS-CoV-2 challenge as suggested by previous reports [51, 52]. Several of the identified differentially regulated genes were of particular interest in the context of adaptive viral T cell immunity (Tables 1 and 2). Several differentially regulated immune response genes laying outside the main window of interest (i.e., Day 5 alone, Day 7 alone, or Day 10 alone) were also identified (Tables 1 and 2 and Supplemental data, Figure S10). For example, on Day 5 in unvaccinated macaques, we found up-regulation of IFIT3 and IL-1RAP. The expression levels of these transcripts have been previously reported to correlate with viral loads in a SARS-CoV-2 rhesus macaque model of COVID-19 disease [5].

**Figure 5.**
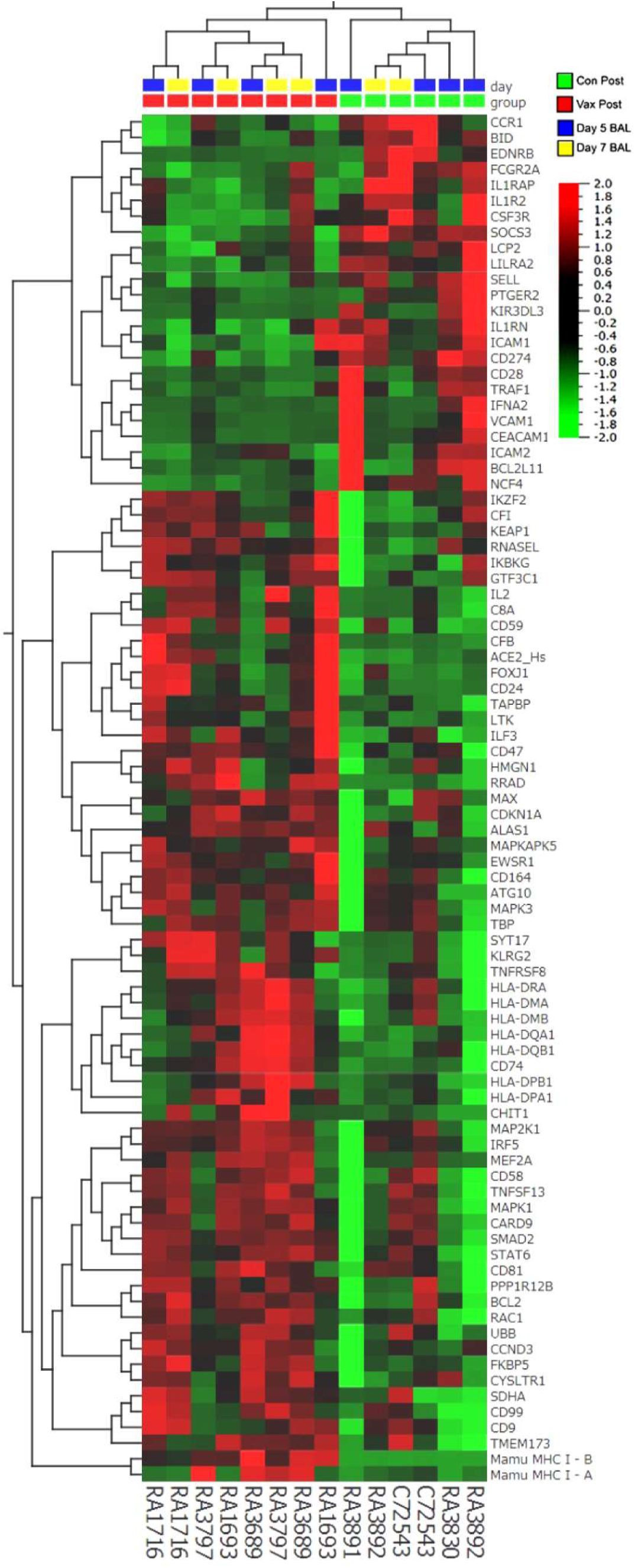
Hierarchical clustering of gene expression in BAL samples collected from control and vaccinated macaques 5- and 7-days post SARS-CoV-2 challenge. Heatmap shows significantly (p<0.05) upregulated (red) genes (63 genes > 3-fold) and down-regulated (green) genes (24 genes < 1/3 fold) from a total of 730 genes analyzed using the NanoString Non-Human Primate Immunology V2 Panel and identifies a set of genes possibly associated with protection from SARS CoV-2 challenge.

**Figure 6.**
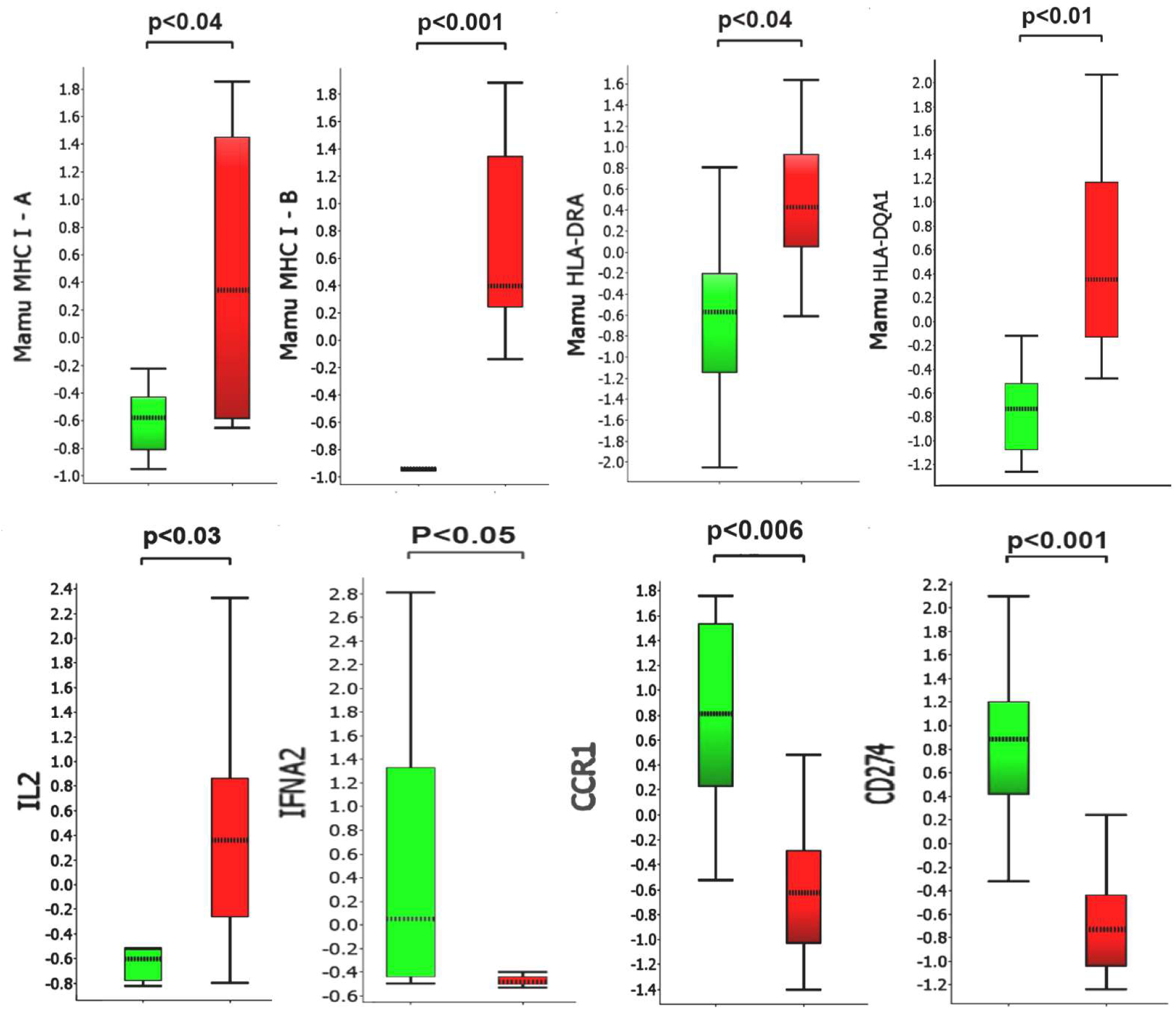
Comparison of selected transcripts up- or down-regulated in collected BAL samples 5-7 days post-challenge. Y-axis values represent fold differences in average scaled counts. Green bars and red bars represent control and vaccinated macaques, respectively. P-values below 0.05 were considered significant.

**Table 1.** Differentially regulated genes in BAL cells obtained from SARS-CoV-2 challenged vaccinated versus control macaques

| Up-regulated Transcripts on Days 5 and 7 Post-Challenge<sup>1</sup> |  |  |  |  |  |
| --- | --- | --- | --- | --- | --- |
| Gene ID | Function | Distribution | Gene ID | Function | Distribution |
| Mamu MHC1 A | Antigen presentation to CD8+ T cells<sup>1</sup> | Low cell type specificity<sup>1</sup> | CD8 | Co-receptor for TCR binding to MHC C1 | T cells |
| Mamu MHC1 B | Antigen presentation to CD8+ T cells | Low cell type specificity | IL2 | Differentiation/maturation of T-cells | CD4+ and CD8+ T cells |
| TAPBP | MHC Class I Antigen presentation | Low cell type specificity | CD81 | Costimulatory signal with CD3 | Low cell type specificity |
| HLA-DRA<sup>2</sup> | Antigen presentation to CD4+ T cells | Professional APC | CD9 | Cell adhesion, recognized by CD81 | Low cell type specificity |
| HLA-DQA1<sup>2</sup> | Antigen presentation to CD4+ T cells | Professional APC | CD59 | Inhibitor of the complement membrane attack complex | Low cell type specificity |
| HLA-DQB1<sup>2</sup> | Antigen presentation to CD4+ T cells | Professional APC | CD24 | Cell adhesion molecule | Eosinophils and B cells |
| CD74 | MHC Class II Antigen presentation | Professional APC | CD47 | High affinity receptor for thrombospondin-1 | Low cell type specificity |
| HLA-DMA | MHC Class II Antigen presentation | Professional APC | CD58 | Ligand of the T-lymphocyte CD2 glycoprotein | Low cell type specificity |
| HLA-DMB | MHC Class II Antigen presentation | Professional APC | CD164 | Facilitates adhesion of CD34+ cells | Low cell type specificity |
| Up-regulated Transcripts on Day 5 or 7 Post-Challenge |  |  | Up-regulated Transcripts on Day 10 Post-Challenge |  |  |
| IL17 B | Proinflammatory cytokine | Low cell type specificity | IL6R | Low affinity receptor for Interleukin 6. | Neutrophil |
| CX3CL1 | Chemotactic for T cells and monocytes | Low cell type specificity | ABL1 | Tyrosine-protein kinase, role cell growth and survival | Low cell type specificity |
| CD99 | Facilitates T cell adhesion | Low cell type specificity | TYK2 | Tyrosine-protein kinase Initiation of type I IFN signaling | Low cell type specificity |

<sup>1</sup>Gene annotations supplied by the human protein atlas [55,56]
<sup>2</sup>Macaque equivalent of human HLA Class II

**Table 2.** Differentially regulated genes in BAL cells obtained from SARS-CoV-2 challenged vaccinated versus control macaques

| Down-regulated transcripts on Days 5 and 7 Post-Challenge<sup>1</sup> |  |  |  |  |  |
| --- | --- | --- | --- | --- | --- |
| Gene ID | Function | Distribution | Gene ID | Function | Distribution |
| IFNA2 | Inhibition of viral replication | Macrophages, eosinophils | CD28 | Provides co-stimulatory signals required for T cell activation. Receptor for CD80 | T cell |
| CCR1 | C-C chemokine receptor, recruitment of immune effector cells | Macrophages | IL1RAP | Coreceptor with IL1R1 in the IL-1 signaling system | Neutrophil |
| CD274 | Ligand of PD-1 (PDL1), inhibits expansion of antigen-specific CD8<sup>+</sup>T cells and CD4<sup>+</sup> helper cells | Monocytes, granulocytes | IL1R2 | Decoy receptor for IL1α, IL1β (IL1B) inhibiting signaling | Macrophage, neutrophils |
| Down-regulated Transcripts on Day 5 or 7 Post-Challenge |  |  | Down-regulated Transcripts on Day 10 Post-Challenge |  |  |
| CD80 | Receptor for CD28 and CTLA-4 on T cells | B cells and monocytes, APCs | HLA-DRA<sup>2</sup> | Antigen presentation to CD4<sup>+</sup> T cells | Professional APC |
| IFNGR2 | $\beta$ chain of the gamma interferon receptor | B cells, APCs and neutrophils | HLA-DMA | Antigen presentation to CD4<sup>+</sup> T cells | Professional APC |
| IL8 | C-X-C chemokine for recruitment of neutrophils | Macrophages, epithelial and endothelial cells | LY96 | Confers responsiveness to LPS | Macrophages |
| IL21 | Regulates proliferation of mature B and T cells in response to activating stimuli | Activated CD4<sup>+</sup> T cells, NKT cells | CTSC | Cathepsin protease | Macrophages |
| DPP4 | Protease upregulated in SARS-CoV-2 [57] Possible viral entry receptor [58] | T cell CD2 | TyroBP | Mediates NK cell activation | Macrophages, monocytes |

<sup>1</sup>Gene annotations supplied by the human protein atlas [55,56]
<sup>2</sup>Macaque equivalent of human HLA Class II

In BAL samples collected on Days 5 and 7, we observed statistically significant up-regulation of MHC Class I genes, MHC Class II and associated accessory genes (CD74 invariant chain, HLA-DM), and T cell markers (CD8 and IL2) in the vaccinated group relative to the unvaccinated control macaques. We also observed statistically significant down-regulation of Interferon alpha 2 (IFNA2), the negative regulator of T cell expansion, PD-L1, the decoy receptor for IL1α and IL1β inhibiting signaling, and FoxJ1, a regulator of Th1 cell activation [53] in the vaccinated group relative to the unvaccinated control macaques.. This pattern suggests enhanced antigen presentation and CD 4/8+ T cell response capacity in BAL cells from vaccinated macaques relative to the controls. In control animals, we observed up-regulation of several genes (CCR1, CSF3R, IFNA2, IL-1RN, IL-1RAP, IL-1R2, and SOXS3) previously reported to be activated during SARS-CoV-2 infection in rhesus macaques [5]. These genes appeared to be down-regulated, relative to controls, in vaccinated macaques.

The vaccine formulation containing synthetic peptide cytotoxic T cell epitopes, CpG, and MPLA adjuvants was delivered primarily via intratracheal instillation. While we did not observe any adverse clinical signs of respiratory distress in vaccinated subjects, we examined BAL cell gene expression for lymphokines and cytokines associated with the observed mixed Th1/Th2 patterns observed in asthma [54] (Supplemental data, Figures S11 and S12). In BAL samples collected on Days 5 and 7, we did not observe significant differences in expression of IL4, IL5, IL6, IL9, IL10. IL13, CXCL10, CCL5, CCL7, CCL22, CX3CL1, or CXCL1. We did observe a general trend of higher expression for many of the cytokines in BAL cell samples collected from vaccinated macaques pre-challenge compared to unvaccinated pre-challenge controls. This effect was generally transient, diminishing by Day 5 post-challenge. This pattern of cytokine and lymphokine expression did not suggest that BAL cells assumed a phenotype associated with a Th2 response.

A major difference between pre-challenge BAL samples was that vaccinated macaques had received prior intratracheal instillation of vaccine formulation containing CpG and MPLA adjuvants. To assess the possible effects and efficacy of our vaccination procedure and formulation, we compared the expression of 60 genes previously reported to be up-regulated, not by antigenic stimulation, but by adjuvants alone [59–63]. We found 30 out of 60 genes examined to be significantly (p<0.05) and differentially (> 2 fold) regulated in BAL cells obtained from vaccinated but unchallenged macaques (Day -7) relative to samples from unchallenged control animals (Day-1) (Supplemental data, Figure S13). This pattern suggests that BAL-associated cells from vaccinated animals were stimulated by the adjuvants in the vaccine formulation. As a further measure of vaccination efficiency, we measured BAL immunoreactivity to the peptides via ELISPOT prior to SARS-CoV-2 challenge (Supplemental data, Figure S14). We observed modest immunoreactivity to one of the six CTL peptides (LL9) in 3 out of the 4 vaccinated pre-challenge macaques. We did not detect immunoreactivity to the peptide antigens using samples of peripheral blood mononuclear cells (data not shown).

## 4. Discussion

In our study, SARS-CoV-2 infection in unvaccinated control macaques progressed similar to previous reports using this model. The post-viral challenge period was clinically characterized by 1) two waves of infectious viral particle recovery in the nasal tissues, 2) lymphopenia on day 1 post-challenge in all animals, and 3) progressive development of pneumonia-like infiltrations visible on chest x-rays as “ground-glass like consolidations”, but few changes associate with human SARS COV-2 infection such as loss of appetite, respiratory distress, vomiting and/or diarrhea. We observed that the kinetics of change in viral loads observed in control rhesus macaques were similar to those previously reported [11]. Additionally, the lung tissue abnormalities revealed by chest radiography were similar in the kinetics of progression and depth to previous reports [6,15,16,20,45–48]. Together, these observations suggest that SARS CoV-2 infection in the Rhesus model is clinically mild, a conclusion confirming some [6,8,11], but not all [64] previous reports. SARS-CoV-2 infection in micro-sphere/adjuvant vaccinated macaques progressed in a pattern different from the unvaccinated controls. Specifically, disease was characterized by a trend toward diminished recovery of infectious viral particles from nasal tissues; enhance recovery of peripheral blood lymphocytes counts, and a significant absence of pneumonia-like infiltrates in the lung. Together, these observations suggest that vaccination conferred some degree of protection against SARS CoV-2 induced disease.

Supporting this clinical conclusion were our studies of the gene expression profiles in serially harvested BAL cells from SARS CoV-2 challenged macaques and immunoreactivity of the BAL cells. In samples collected from vaccinated (but pre-viral challenge) macaque BAL cells, we found distinct changes in gene expression associated with the use of TLR 4 and 9 agonists, including up-regulation of CSF1 [61], IRF7 [62], and IL10[60] suggesting effective delivery of the adjuvant portion of the vaccine formulation. When we examined HLA Class I restricted immunoreactivity of the pre-challenge but vaccinated macaque BAL cells towards the vaccinating peptides, we found modest to low reactivity towards one of the peptides (LLLDRLNQL) in three of four NHP subjects vaccinated.

Following vaccination and viral challenge, we found evidence of up-regulation of both Mamu MHC Class I and Class II genes in macaque BAL cells relative to unvaccinated, viral challenged macaques. Upregulation of HLA Class I and Class II molecules in peripheral blood mononuclear cells has been reported following the measles /mumps/rubella (MMR) vaccine in MMR naïve individuals [65]. Increased signatures of M1-type macrophage APC transcripts in the BAL of SARS-CoV-2 infected rhesus macaques has been previously reported [5]; however our finding of upregulated expression of MHC Class II genes, a hallmark of professional APCs, appears to be unique to the BAL of our vaccinated macaques. Likewise, we found increased up-regulation of the IL2 genes in vaccinated relative to unvaccinated macaques. A similar finding has been reported in humans where higher IL 2 levels distinguish mild/asymptomatic forms of COVID-19 disease from the moderate/severe forms [66]. Following viral challenge in our study (specifically, Day 5 to 7), we found down-regulation of IFNα2 genes in BAL cells recovered from vaccinated subjects relative to their unvaccinated controls. The observation perhaps reflects the decreased viral loads found in vaccinated NHP subjects [5].

While antibodies are a critical component of the protective humoral immune response to pathogens, antibodies that promote disease have been described and categorized as ADE [67] or VAERD, such as that described in MERS-CoV or RSV patients [68–70]. Since our vaccine only delivered low molecular weight nonameric synthetic peptides, unconjugated to any carrier, we did not expect to generate a significant anti-peptide humoral response. Rather, during the development and testing of the microsphere synthetic peptide COVID 19 vaccine, we were mindful of evoking Th2 biased immune responses, particularly those that occur in the absence of Th1 responses or appropriate T regulatory cell responses [71]. The gene expression patterns of BAL cells obtained from the vaccinated macaques in our study suggested that we did not provoke an unbalanced Th2 response by vaccination.

We note several weaknesses in our approach to demonstrate the efficacy of this experimental vaccine: 1) SARS-CoV-2 infection in Rhesus macaques resulted in only mild disease, which appears to resolve by day 10-14 post-infection. Having noted this in previous reports on the SARS-CoV-2 /rhesus model, we tried to induce more severe forms of infection using higher doses of infectious particles than previous reports. Based on clinical signs, we found little effect of the increased dose of the virus on Rhesus SARS-CoV-2 disease severity. 2) Because of the limited amount of cells recovered from the BAL, we were where unable to perform confirmatory quantitative RT-PCR analysis of the unique gene expression patterns found in vaccinated versus control BAL. Nevertheless, we found that many of our observations have been previously reported from studies in similar models, 3) Not all the macaque chest radiographs were performed on the same day in control versus vaccinated subjects, a reflection of the logistic difficulties in working under BSL ¾ conditions. We are confident, however, we have captured the chest radiographic abnormalities induced by SAR-CoV-2 infection of macaques (i.e. conspicuous consolidations and infiltrations prevalent in the caudal lobe of the right lung), and adequately shown their absence in vaccinated viral challenged macaques, 4) we only included one cynomolgus macaque in our control group. This was primarily due to macaque availability at the time the study was conducted. However, this appears to have been both a strength [64] and weakness of the experimental design, as we found similar lung pathology in the cynomolgus macaque as well as similar patterns of BAL gene expression as the controls Rhesus subjects [72], and 5) we did not study the effects of adjuvant alone on SARS-Cov-2 infection in the Rhesus macaques. Previous experience with this microsphere CTL vaccine platform in a murine Ebola virus model has shown that adjuvant alone was not sufficient to confer protection against lethal virus challenge. Protection was conferred only when the corresponding synthetic CTL peptide epitopes were delivered in the microsphere [4].

We believe this report is the first demonstration of efficacy in a preclinical NHP model of SARS CoV-2 infection of a synthetic peptide-based vaccine based on known and persistently immunogenic HLA Class I bound CTL peptide epitopes of SARS nucleoprotein [28]. The SARS-CoV-2 nucleoprotein genomic sequences have shown significantly reduced mutations rates compared to spike protein. As such, it may represent an additional target for vaccination, perhaps in the context of a booster vaccine used following SARS-CoV-2 spike protein vaccines based on recombinant protein, mRNA, or adenoviral vectors [73–75]. The ready ability to change the sequence of the synthetic peptide HLA Class I restricted CTL epitopes used in the system is an attractive feature, given the observed rates of mutation in SARS-CoV-2 as it spreads in the human population in the future. A second potentially attractive feature of this vaccine approach is that it can be delivered by aerosolization to the respiratory mucosa, a route previously demonstrated to generate efficiently lung-dwelling tissue-resident memory T cells [76, 77].

## 5. Conclusions

We demonstrate that Rhesus macaques receiving the microsphere vaccine formulation prior to viral challenge are protected from pneumonia-like lung abnormalities that characterize SARS CoV-2 infection in unvaccinated control macaques. Analysis of gene expression of cells obtained from bronchiolar lavage shows unique signatures consistent with the hypothesis that vaccination with this platform induces a protective T cell response in viral challenged macaques.

## 6. Conflict of Interest / Competing Interests Disclosures and Contributions

R.R., T.Bl., S.B., S.C., R.C., T.H., L.W., P.L., and C.H. are employees of Flow Pharma, Inc. compensated in cash and stock, and are named inventors on various issued and pending patents assigned to Flow Pharma. Some of these patents pending are directly related to the study presented here. P.H. is a member of Flow Pharma’s Scientific Advisory Board. T.Be. is a Flow Pharma stockholder. The remaining authors declare that the research was conducted in the absence of any commercial or financial relationships that could be construed as a potential conflict of interest. All authors made substantial contributions to: (1) the conception and design of the study (R.R., S.B., S.C.,T.Br., J.C., P.H., C.H., T.Be.), or acquisition of data (T.Br. C.M. J.C., C.H.), or analysis and interpretation of data (R.R., P.H., T.Br., J.C.), (2) drafting the article or revising it critically for important intellectual content (P.H., T.Br. S.B., T.H., T.Bl., P.L., R.R.) (3) and all authors have approved the final version of the submitted manuscript.

## 7. Acknowledgments

The authors wish to thank Dr. Antonella Maffei for her critical review of the manuscript.

## 9. Data Availability Statement

The original DICOM files (> 20 MB/file) are available upon request.

## 10. Appendix

none

## Supplemental Materials

**Supplemental Data, Figure S1.**
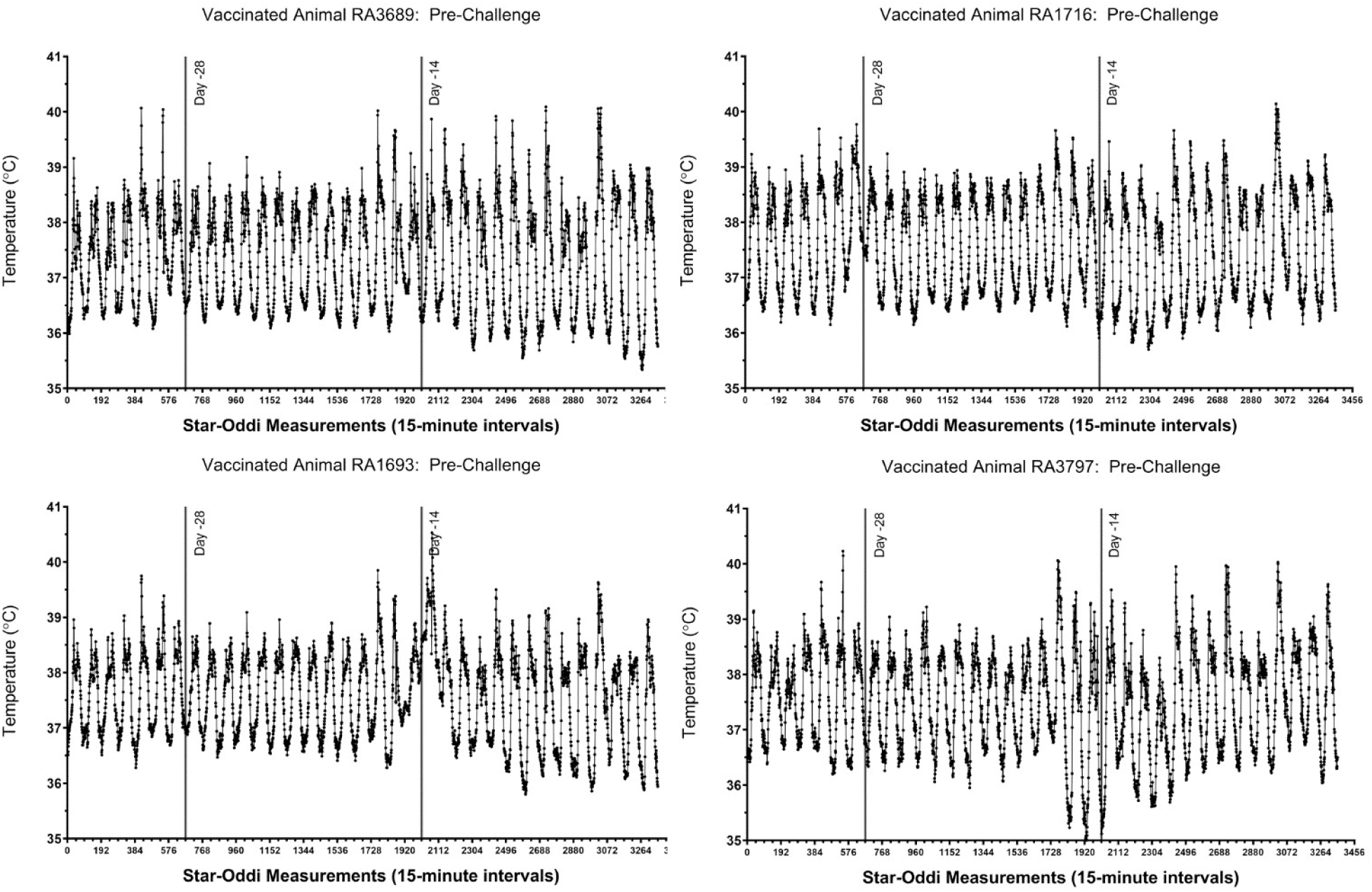
Core body temperature alterations in vaccinated macaques prior to SARS-CoV-2 challenge. For each animal, 35 days of pre-challenge temperature measurements are shown. Each tick on the x-axis represents 12 hours or 48 individual logger measurements.

**Supplemental Data, Figure S2.**
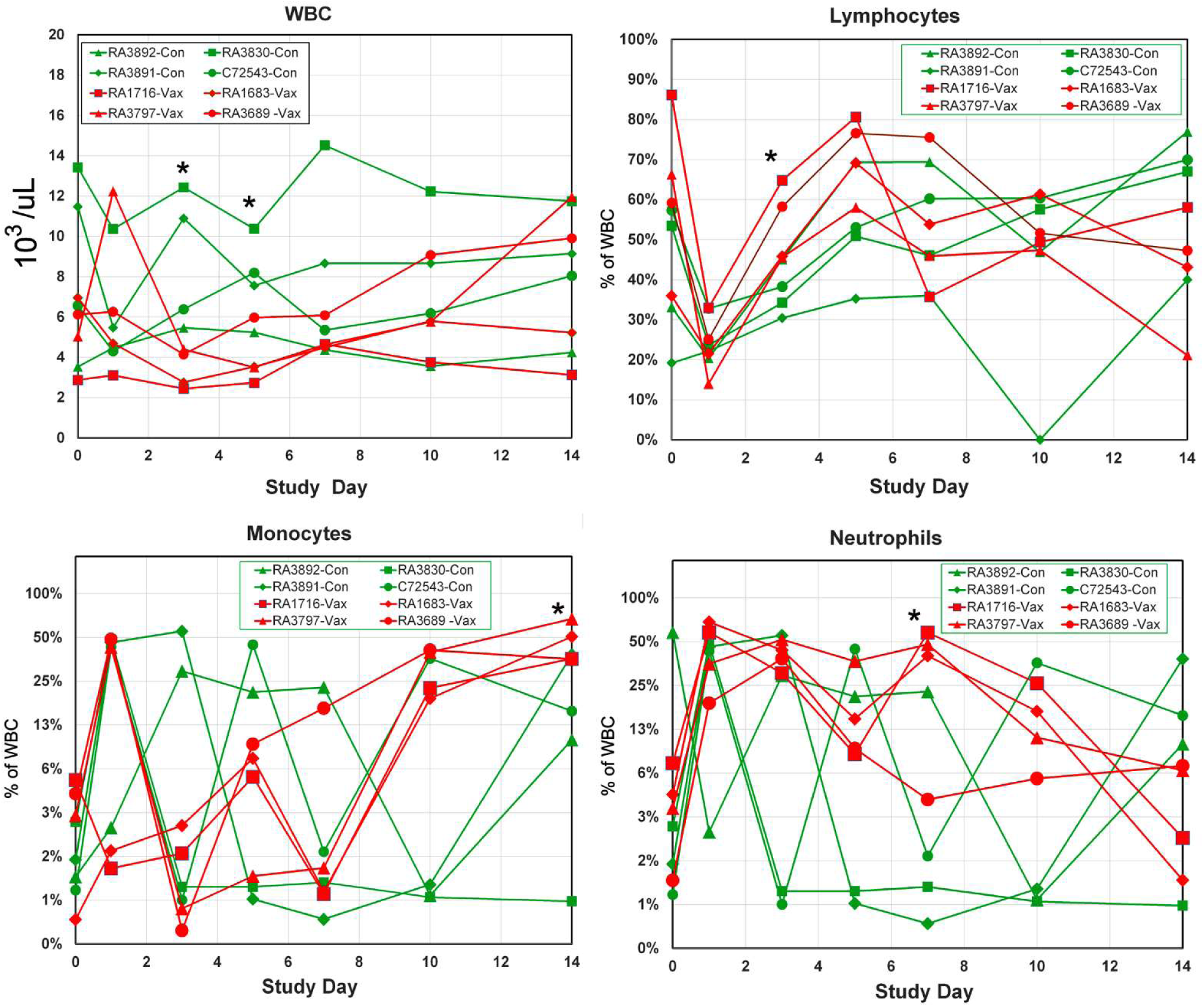
Hematological analysis in control and vaccinated macaques challenged with SARS-CoV-2. The counts of white blood cells (WBC) (upper left panel), the percent of lymphocytes in WBC (upper right panel), the percent of monocytes in the WBC (lower left panel), and the percent of neutrophils in the WBC (lower right panel) were analyzed. An asterisk indicates a statistically significant difference (p< 0.05) between control and vaccinated macaques by Students t-test.

**Supplemental Data, Figure S3.**
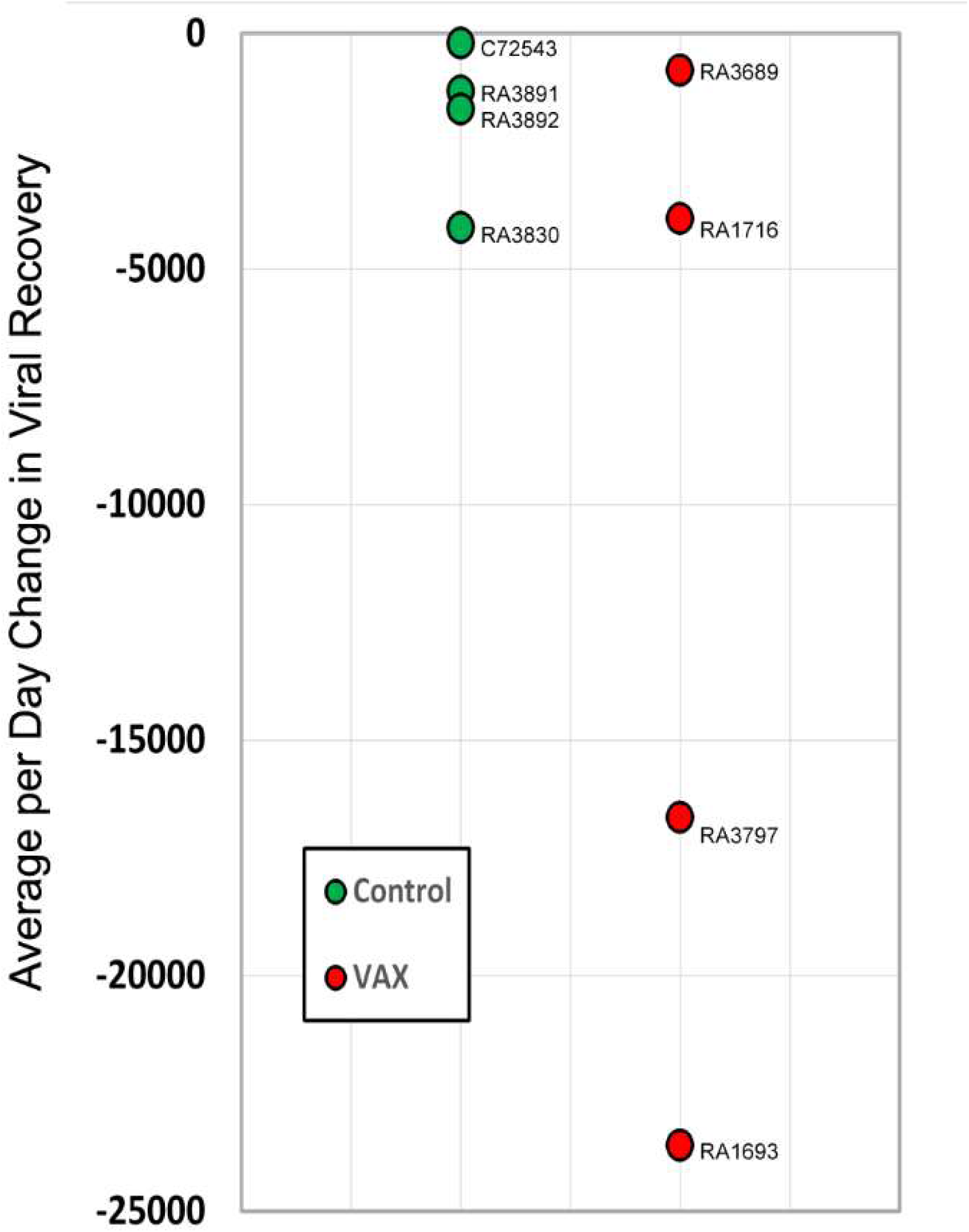
Viral clearance rates in control and vaccinated SARS CoV-2 challenged macaques. From the total viral loads measurements in nasal swab samples from SARS-CoV-2 challenged macaques, the daily viral clearance rates (i.e. TCID _50_ /mL _day n-1_ minus TCID _50_ /mL _day n_) were calculated and averaged over a nine-day period. Red symbols are vaccinated macaques subjects, control unvaccinated macaque subjects are shown in green symbols.

**Supplemental Data, Figure S4.**
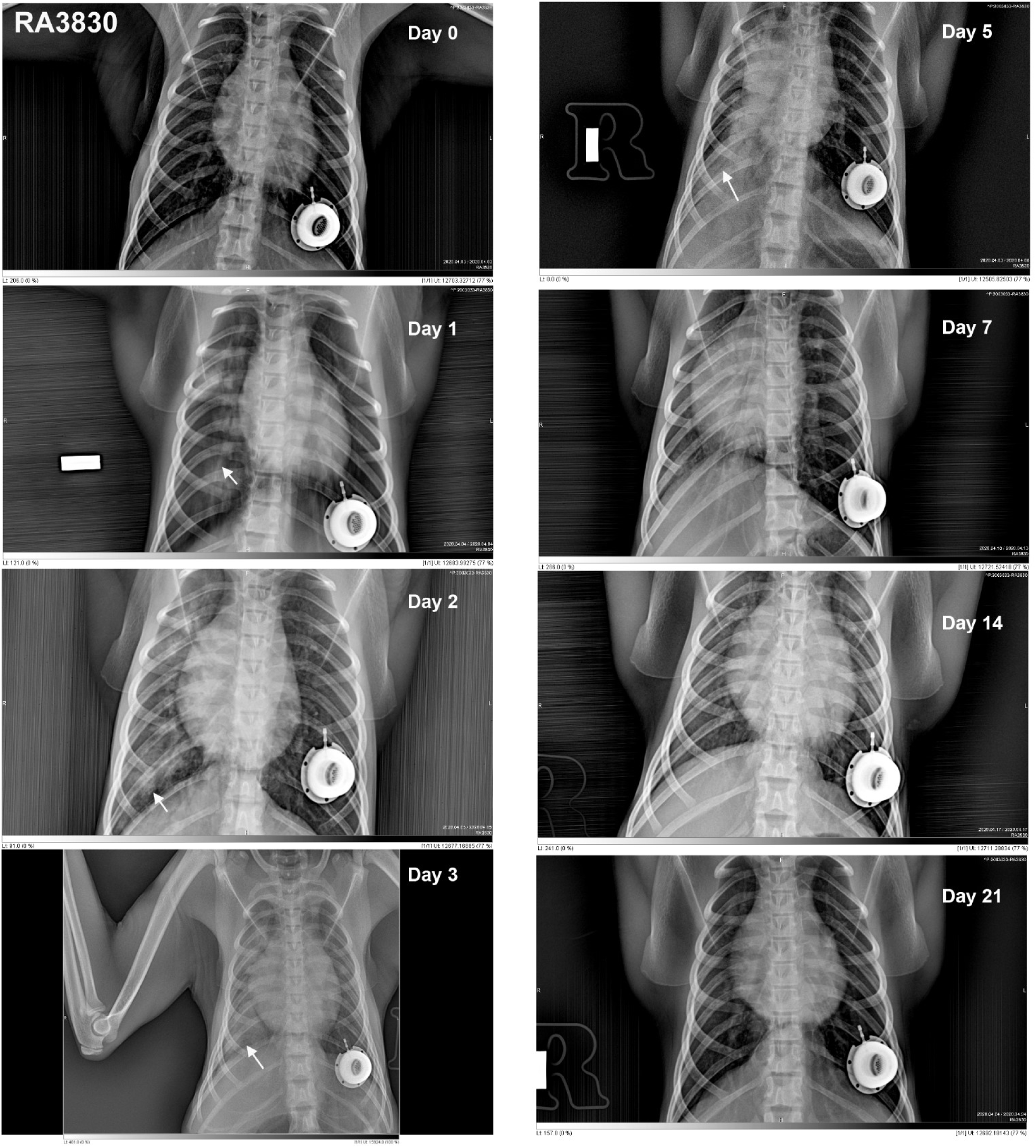
Chest radiographs of control rhesus macaque RA3830 following SARS-CoV-2 challenge. As shown, this animal demonstrated a progression of pulmonary infiltrates during the acute period (Days 2-5) of disease post-challenge which resolved by study termination (Day 21). White arrows indicate areas of mild to moderate pulmonary infiltrates seen as ground glass consolidations.

**Supplemental Data, Figure S5.**
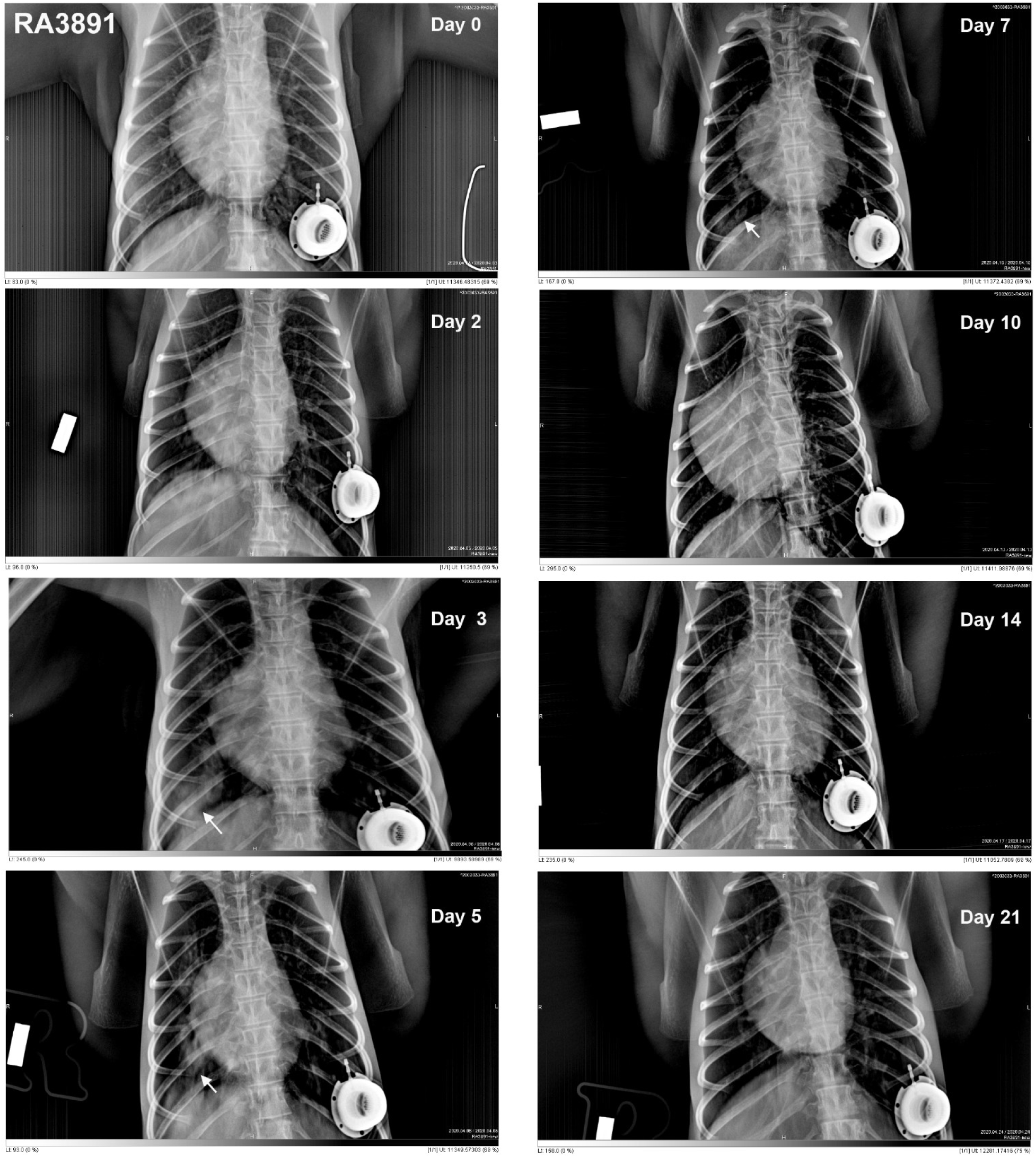
Chest radiographs of control rhesus macaque RA3891 following SARS-CoV-2 challenge. As shown, this animal demonstrated a progression of pulmonary infiltrates during the acute period (Days 3-7) of disease post-challenge which resolved by study termination (Day 21). White arrows indicate areas of mild to moderate pulmonary infiltrates seen as ground glass consolidations.

**Supplemental Data, Figure S6.**
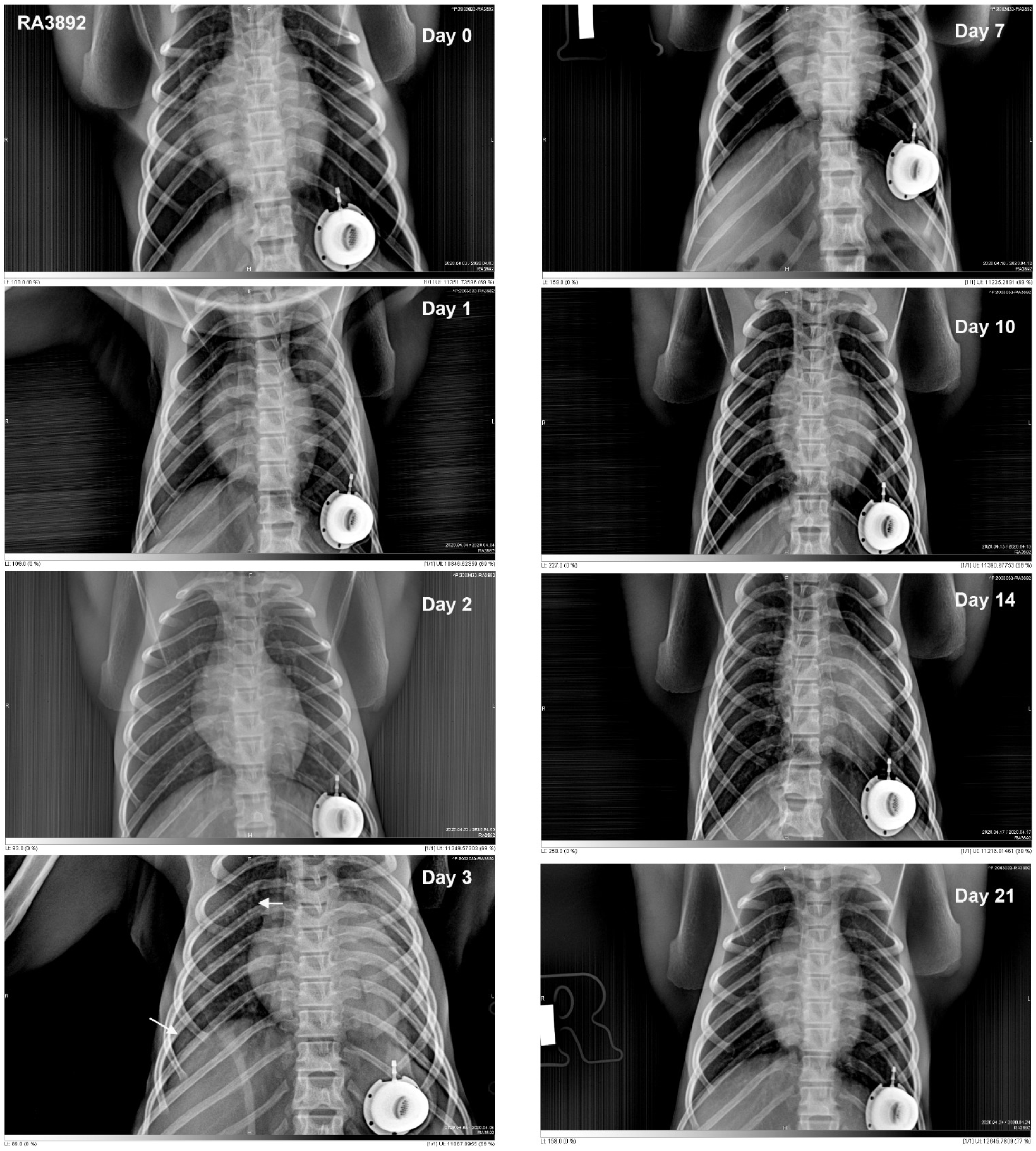
Chest radiographs of control rhesus macaque RA3892 following SARS-CoV-2 challenge. As shown, this animal demonstrated a progression of pulmonary infiltrates during the acute period (Day 3) of disease post-challenge which resolved by study termination (Day 21). White arrows indicate areas of mild to moderate pulmonary infiltrates seen as ground glass consolidations.

**Supplemental Data, Figure S7.**
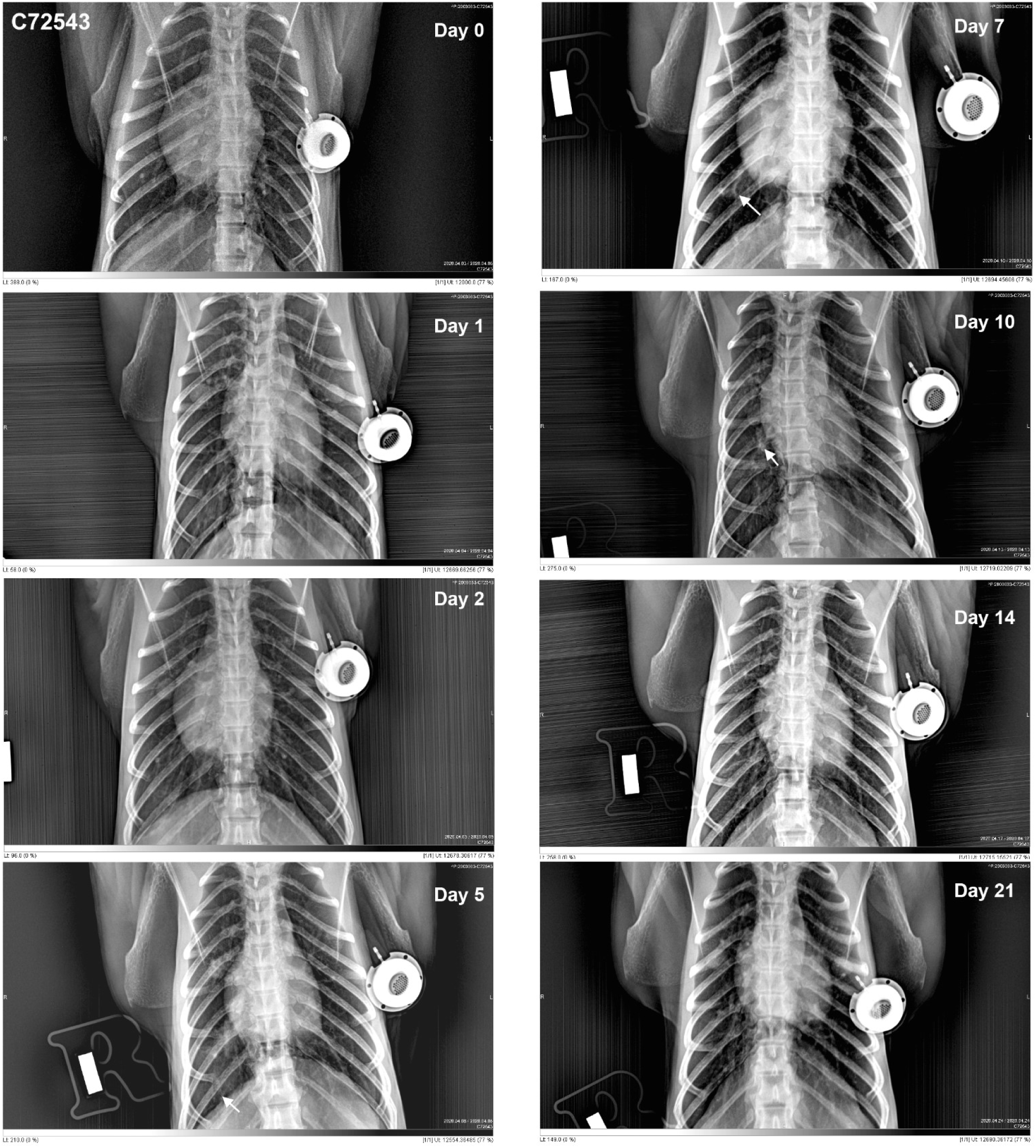
Chest radiographs of control cynomolgus macaque C72543 following SARS-CoV-2 challenge. As shown, this animal demonstrated a progression of pulmonary infiltrates 5-10 days post-challenge which resolved by study termination (Day 21). White arrows indicate areas of mild to moderate pulmonary infiltrates seen as ground glass consolidations.

**Supplemental Data, Figure S8.**
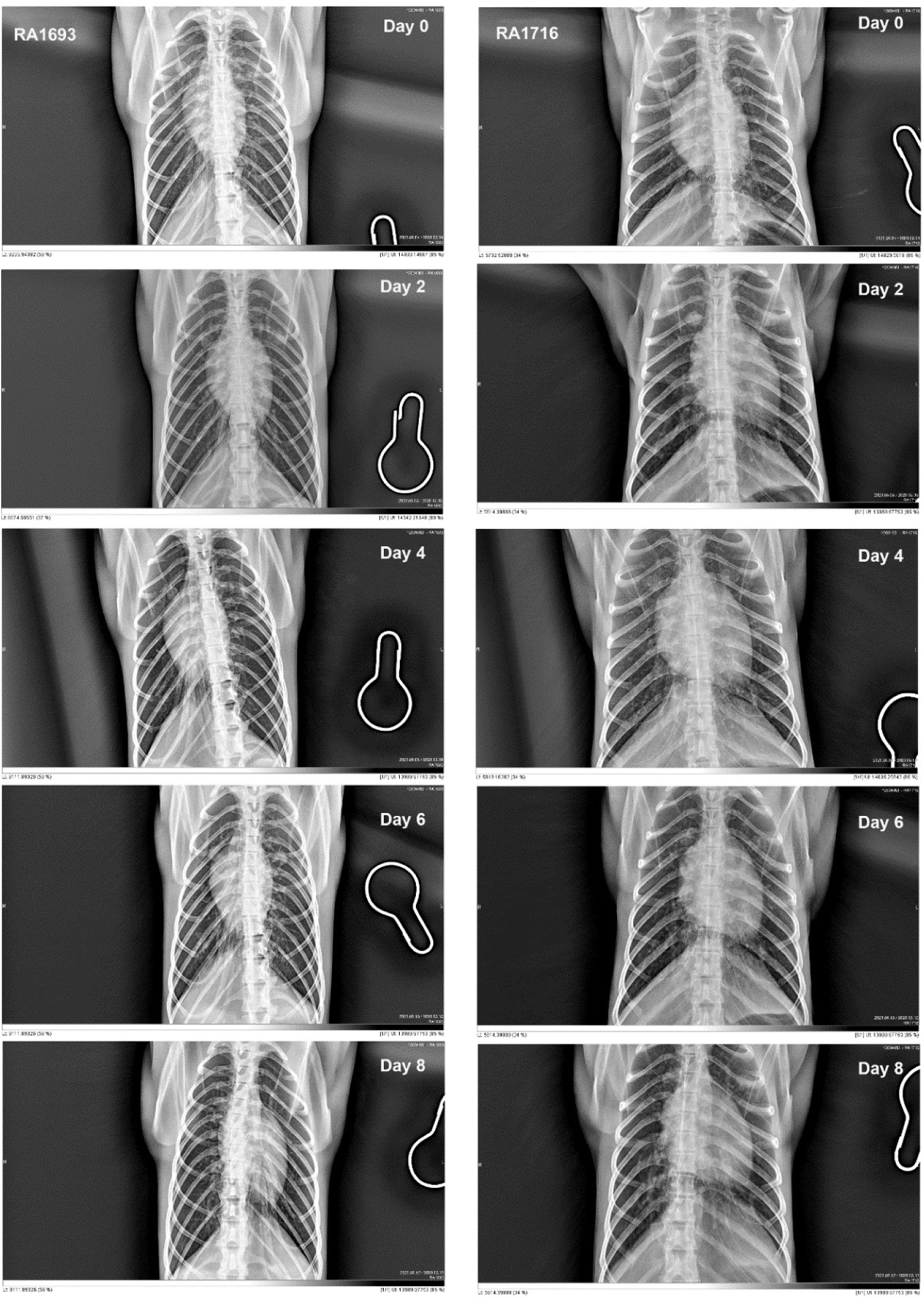
Chest radiographs of vaccinated rhesus macaques RA1693 and RA1716 following SARS-CoV-2 challenge. With the exception of increased reticulation relative to baseline, few abnormalities were observed in collected radiograph images. Note the absence of infiltrates or consolidation typically seen in the unvaccinated control population.

**Supplemental Data, Figure S9.**
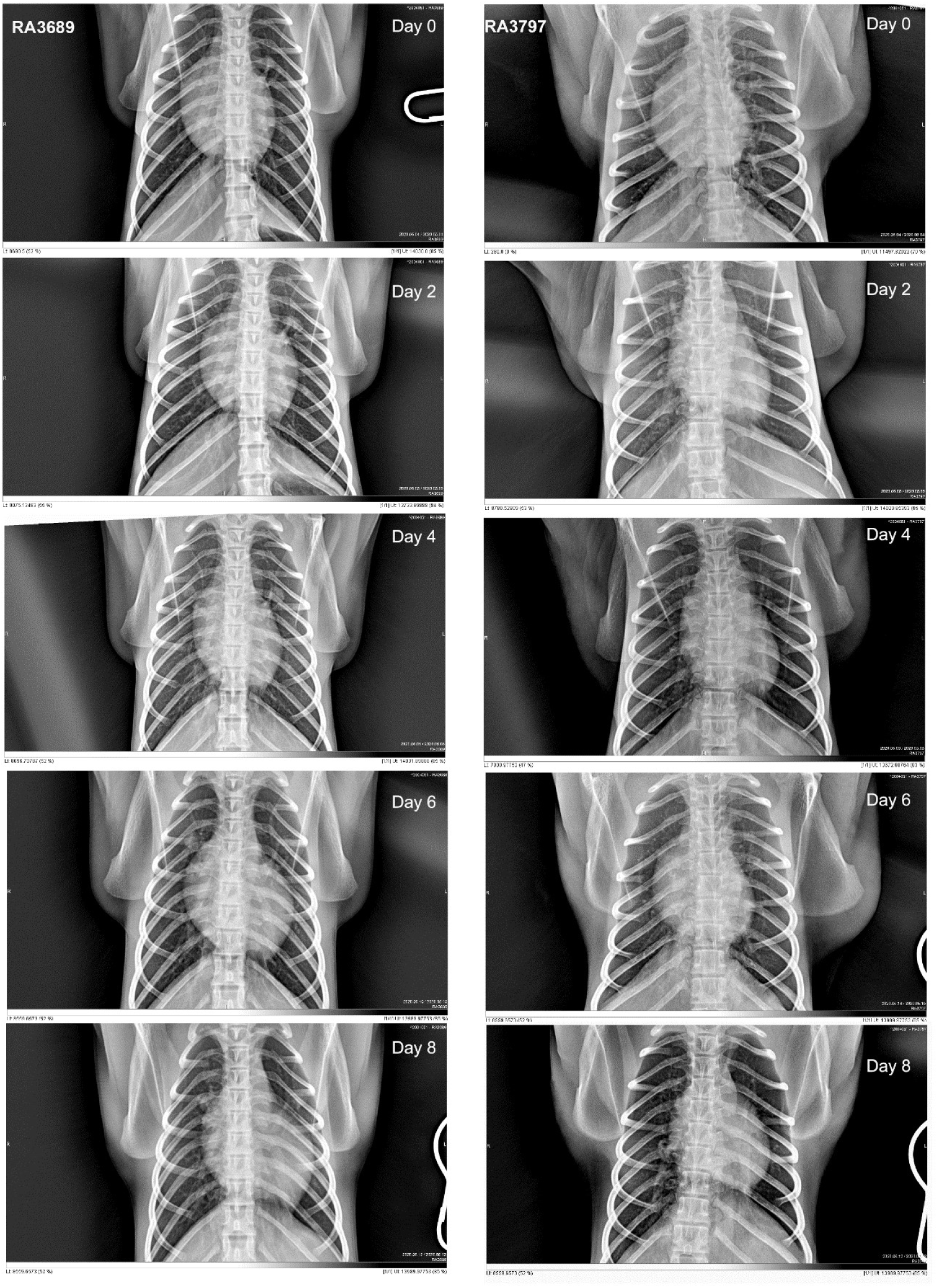
Chest radiographs of vaccinated rhesus macaques RA3689 and RA3797 following SARS-CoV-2 challenge. Radiographs are unremarkable, other than showing increased reticulation relative to baseline, appearing on days 2 through 4, clearing on later imaging. In particular, note lack of focal infiltrates or consolidations.

**Supplemental Data, Figure S10.**
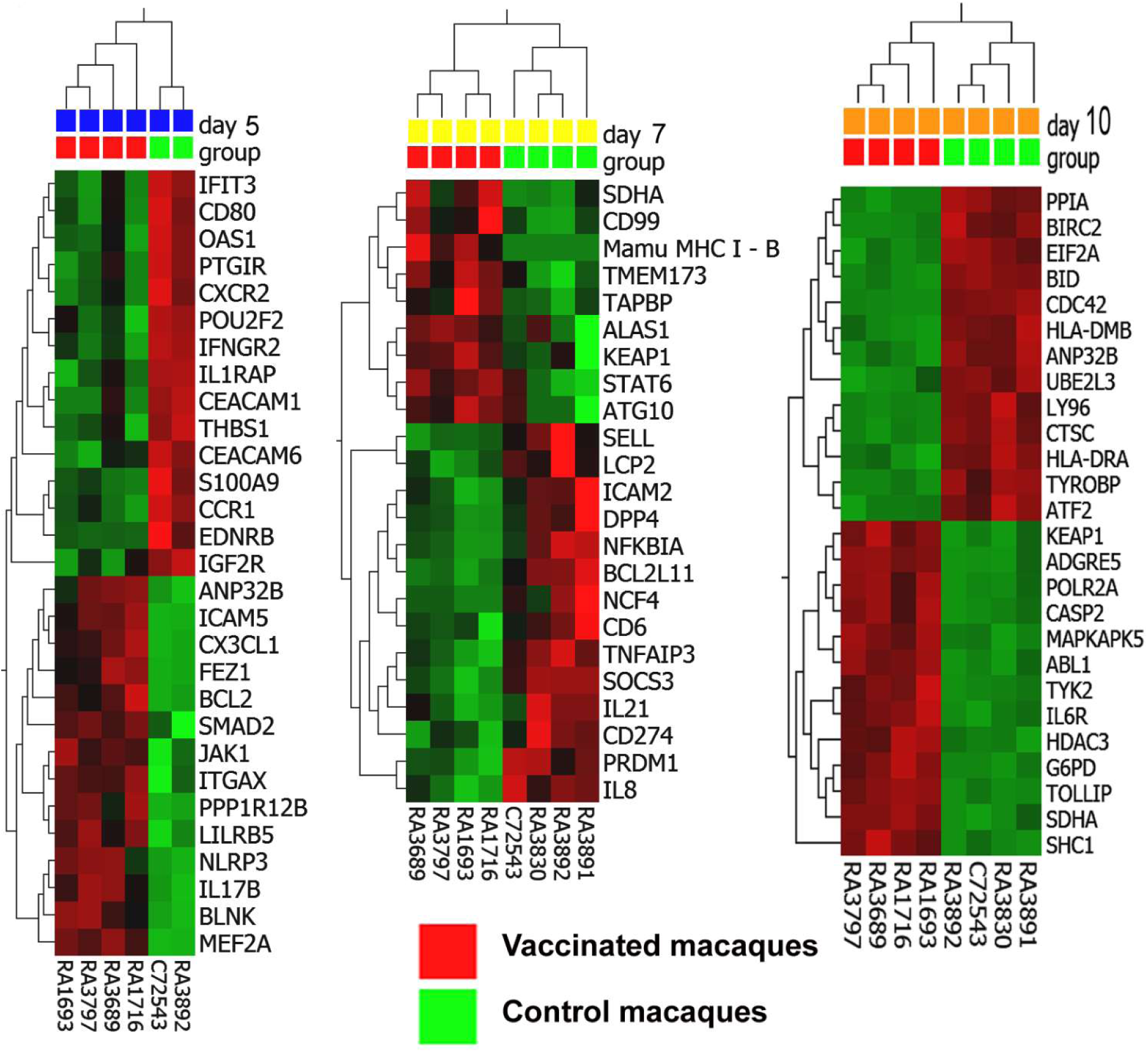
Hierarchical clustering of gene expression in BAL samples collected from control and vaccinated macaques on Day 5 (left panel), Day 7 (middle panel), and Day 10 (right panel). 5- and 7-days post SARS-CoV-2 challenge. Heatmap shows significantly (p<0.05) up-regulated (red) transcripts and down-regulated (green) transcripts from a total of 730 genes analyzed using the NanoString Non-Human Primate Immunology V2 Panel.

**Supplemental Data, Figure S11.**
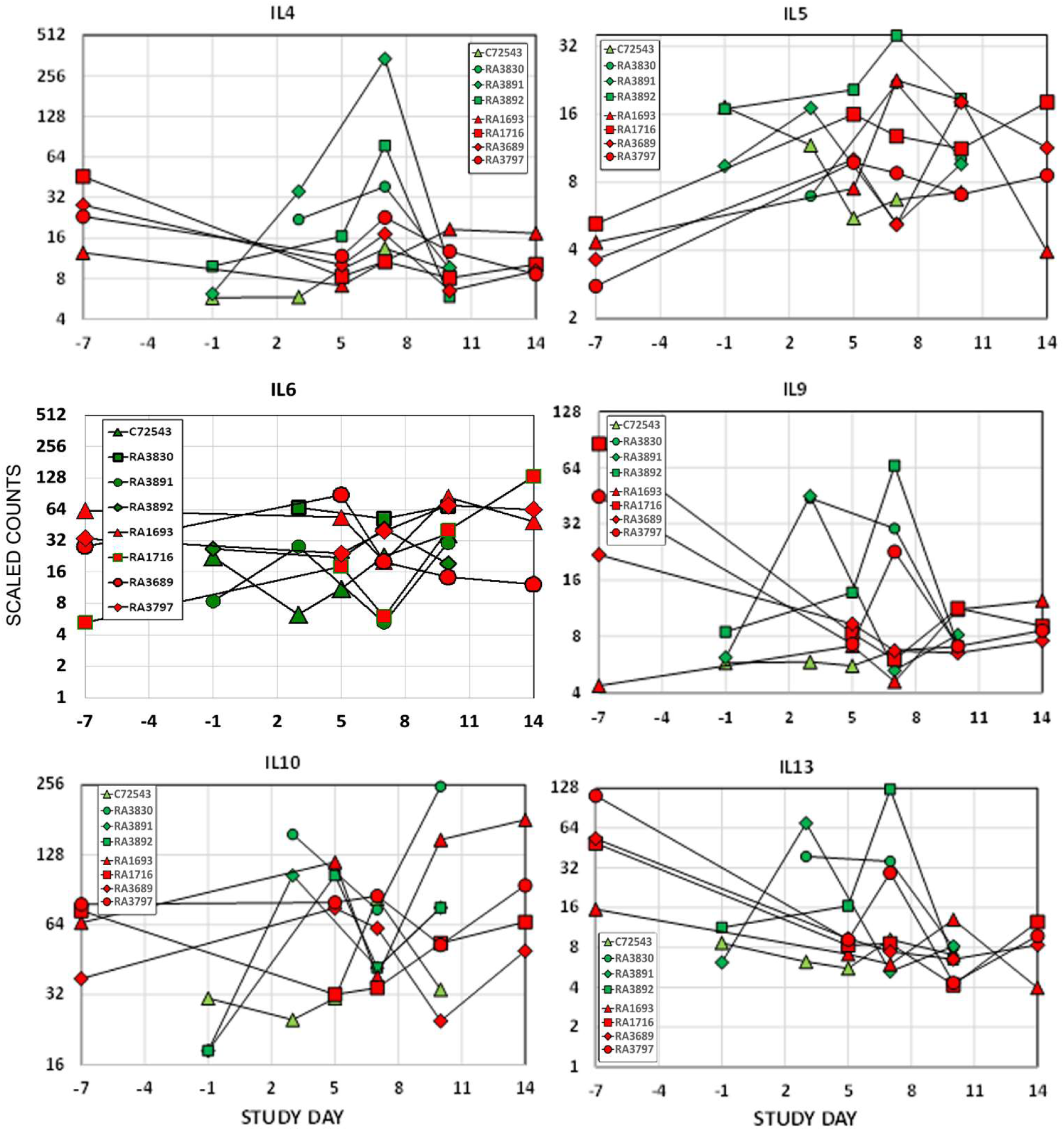
Intratracheal vaccination with the adjuvanted micro-sphere peptide vaccine did not promote the expression of Th_2_ type interleukin cytokine transcripts in collected BAL samples relative to levels measured in control macaques. BAL cell gene expression (shown as scaled counts on the y-axis) of cytokines associated with Th_2_ T cell responses were plotted by study day (x-axis) for each animal prior to and following the SARS-CoV-2 challenge. No statistically significant differences between control and vaccinated macaques were found.

**Supplemental Data, Figure S12.**
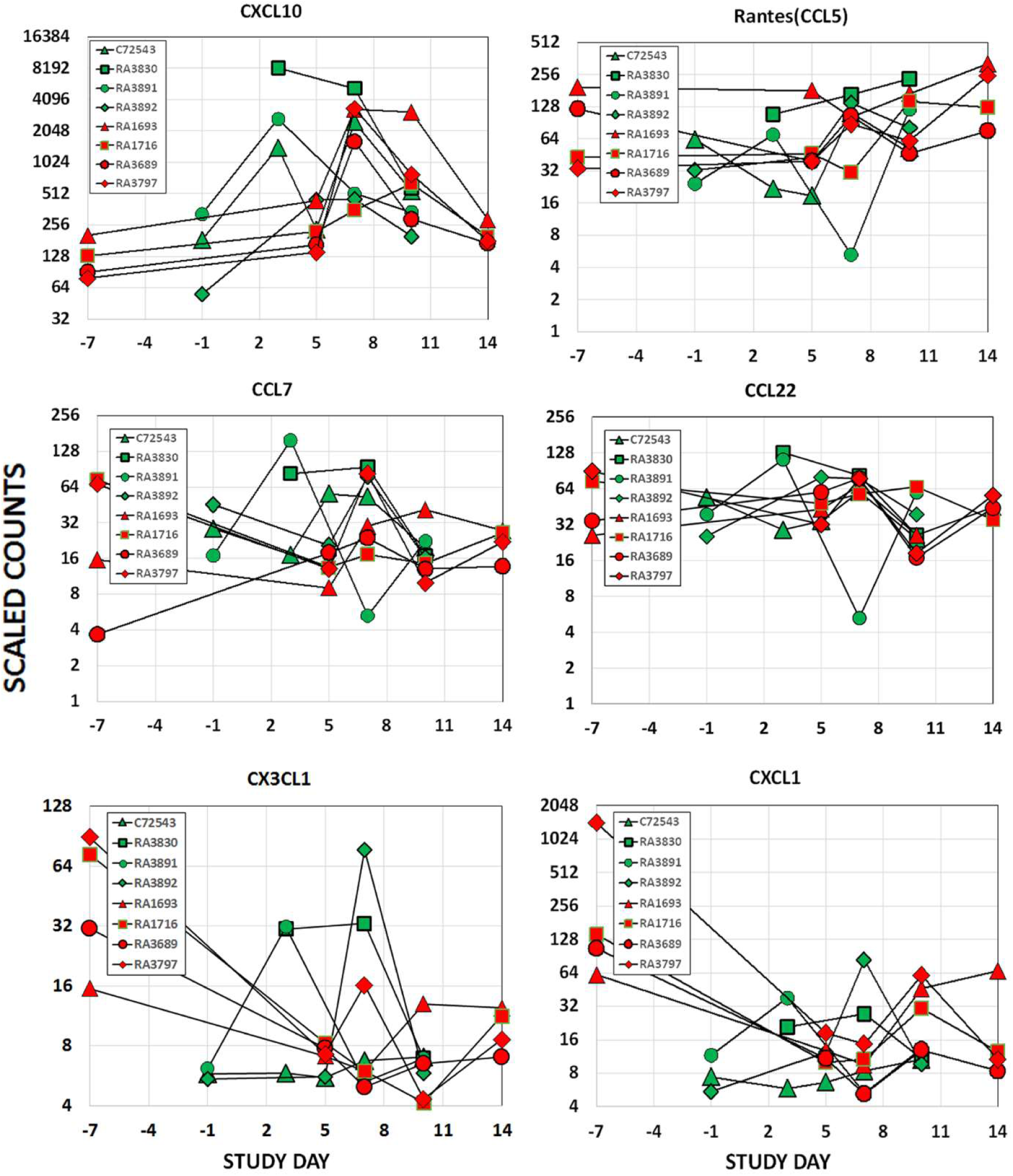
Intratracheal vaccination with the adjuvanted microsphere peptide vaccine did not promote the expression of Th_2_ type interleukin cytokine transcripts in collected BAL samples relative to levels measured in control macaques. BAL cell gene expression (shown as scaled counts on the y-axis) of cytokines associated with Th_2_ T cell responses were plotted by study day (x-axis) for each animal prior to and following the SARS-CoV-2 challenge. No statistically significant differences between control and vaccinated macaques were found.

**Supplemental Data, Figure S13.**
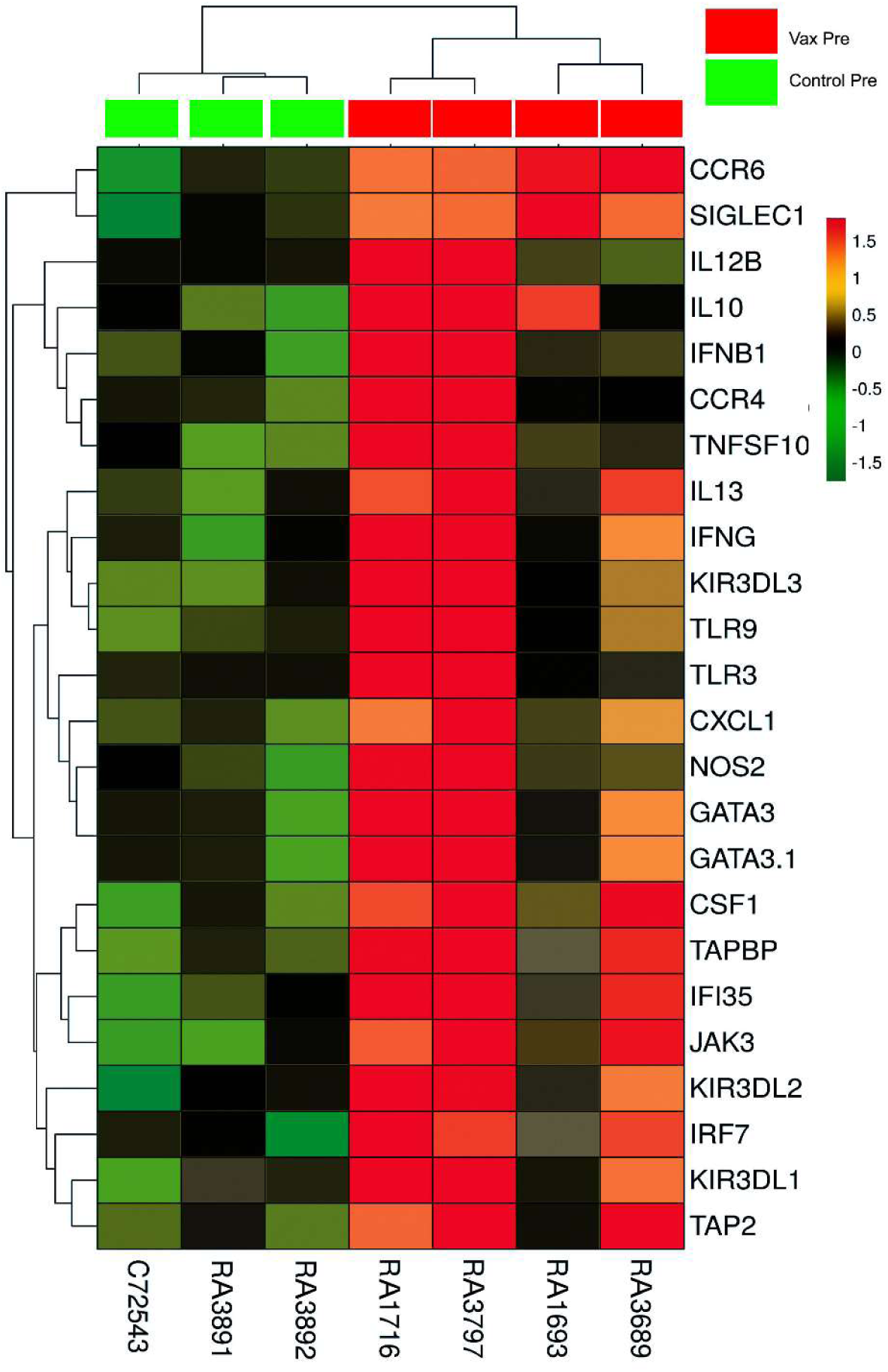
Hierarchical clustering of adjuvant-related transcript expression in BAL samples collected from control and vaccinated macaques prior to virus challenge (indicated as Control Pre and Vax Pre). Gene expression analysis identified differentially regulated genes in BAL samples obtained on Day -1 from control animals and Day -7 from vaccinated macaques. Heatmap shows significant (p <0.05) differential expression of a series of genes that had previously been identified (see main text) as being regulated by adjuvants alone. Up-regulated (red); down-regulated (green). Macaque RA 3830 was not sampled on day - 1.

**Supplemental Data, Figure S14.**
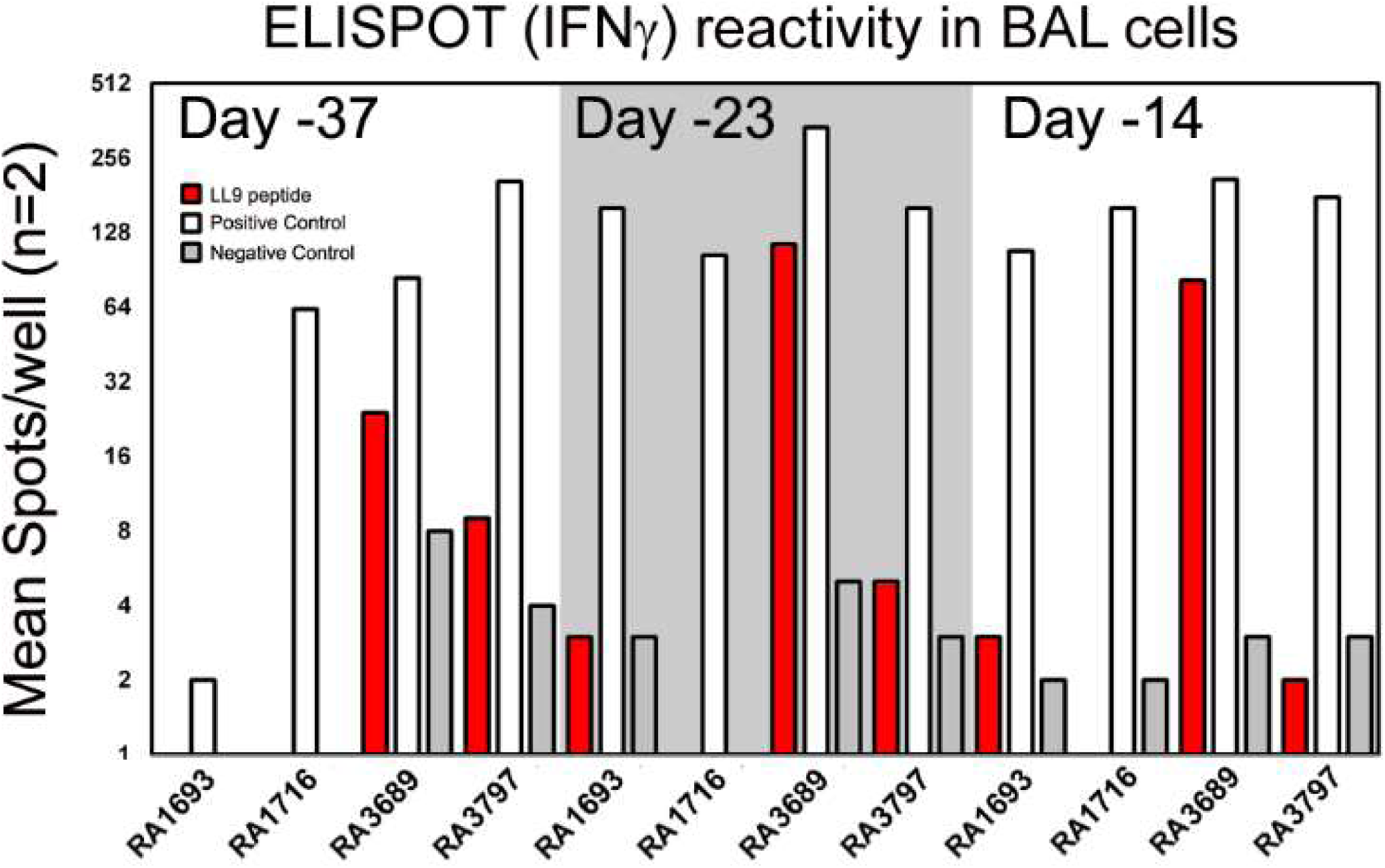
Immunoreactivity of BAL-associated cells from vaccinated macaques to immunizing peptides prior to SARS-CoV-2 challenge. Dates shown are assay date rather than sampling date. Concanavlin A was used as a positive control.

**Table S1A.**
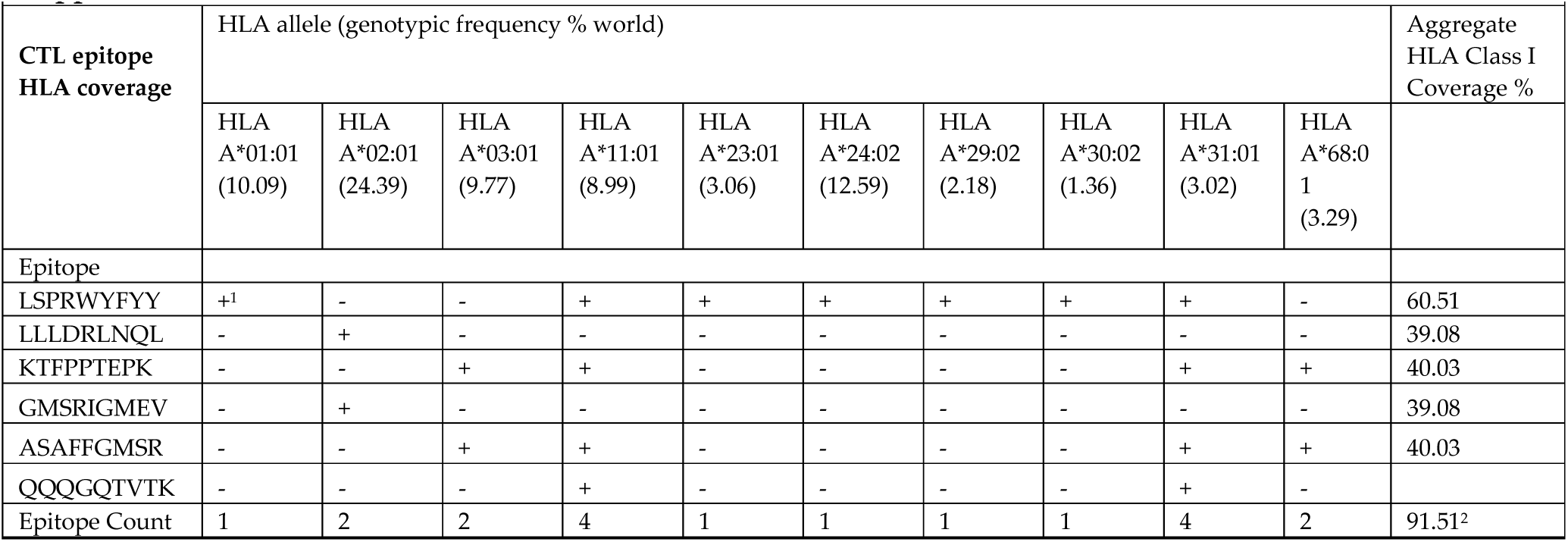
Notes: 1. + Indicates positive in-vitro assays for MHC binding and/or T-cell recognition [36]. 2. Calculated as previously described[78]

**Table S1B.**
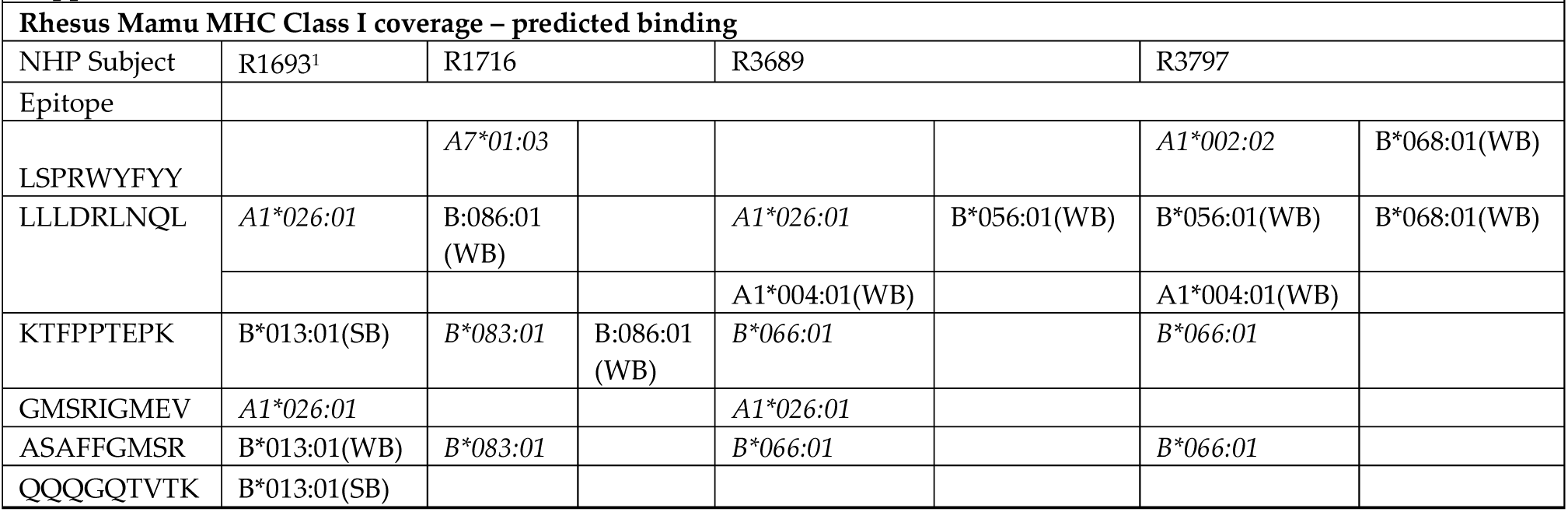
Notes: 1. A typical rhesus MHC haplotype may contain two or three expressed Mamu-A genes, and up to nineteen distinct Mamu-B-like loci[79], 2. Mamu MHC in italics are predicted to bind based on HLA homology and in-vitro analysis [40]. 3. Mamu MHC in regular font are predicted to bind based on NetMHCpan 4.1, with a weak binder (WB) at top 2% percentile rank and strong binder (SB) at top 0.5% precentile rank.

